# REFEE Shapes ZGA Through Post-Transcriptional Regulation of Repetitive RNAs in Mouse Embryos

**DOI:** 10.1101/2025.08.21.671216

**Authors:** Mie Kobayashi-Ishihara, Akihiko Sakashita, Youjia Guo, Hirotsugu Ishizu, Hidetoshi Hasuwa, Ten D. Li, Toshiya Nakahara, Tomohiro Kitano, Hidetaka Kosako, Kensaku Murano, Haruhiko Siomi

**Affiliations:** Department of Molecular Biology, Keio University School of Medicine, Tokyo, Japan; The University of Tokyo Pandemic Preparedness, Infection and Advanced Research Center (UTOPIA), The University of Tokyo, Tokyo, Japan; Laboratory for Developmental Genome Plasticity, RIKEN BioResource Research Center, Ibaraki, Japan; Department of Biomedical Sciences, School of Veterinary Medicine, University of Pennsylvania, Pennsylvania, USA; Laboratory Animal Center, Keio University School of Medicine, Tokyo, Japan; Division of Cell Signaling, Institute of Advanced Medical Sciences, Tokushima University, Tokushima, Japan; Human Biology Microbiome Quantum Research Center (WPI-Bio2Q), Keio University, Tokyo, Japan; Center for Next Generation In Vivo Research, Chiba University, Chiba, Japan

**Author notes:** To whom correspondence should be addressed (HS), (MKI).

## Abstract

During early mammalian embryonic development, repetitive elements such as transposons and duplicated genes are transcriptionally active and play key roles in embryogenesis. MERVL, an endogenous retrovirus, is essential for shaping the early transcriptome, particularly during the transition from totipotency to pluripotency. However, the function of many repetitive elements remains poorly understood. Here, we add a new repetitive player, REFEE, that regulates repetitive RNAs and contributes to totipotency regulation through post-transcriptional mechanisms. REFEE was initially identified as a paralogue of *Alyref*, a nucleocytoplasmic RNA exporter, transiently expressed during the 2-cell stage of mouse embryogenesis. REFEE preferentially binds and stabilizes MERVL RNA, promoting its export. REFEE knockdown embryos fail to complete zygotic genome activation (ZGA), accompanied by a significant reduction in MERVL RNA levels. REFEE primarily influences transcript levels of ZGA genes, upregulating or downregulating them in a manner correlated with their genomic proximity to MERVL. Proteomic analysis further revealed that REFEE knockdown also decreases protein levels of its RNA targets, indicating an additional layer of post-transcriptional regulation beyond RNA stability. Our findings identify REFEE as an essential post-transcriptional regulator of early embryonic development and highlight the functional interplay between repetitive elements in shaping the zygotic transcriptome.

## Introduction

The earliest stages of mammalian embryonic development exhibit a distinct transcriptomic landscape, markedly different from later life stages. This early transcriptome comprises maternal RNAs inherited from the oocyte, as well as transcripts derived from transposable elements and multicopy genes. The expression of these repetitive elements, mainly included in the so-called “2-cell (2C) transient genes” or “2C genes” in mice^1^, has traditionally been considered a promiscuous feature of early embryos, primarily attributed to the loosely organized and permissive chromatin present immediately after fertilization^2,3^. This chromatin structure undergoes progressive reorganization during zygotic genome activation (ZGA), a crucial process that drives the stepwise onset of transcription in the zygote while facilitating the degradation of maternal transcripts. Intriguingly, expressions of these 2C-stage transcripts seem to coincide with the totipotent plasticity of the preimplantation embryo^1^. However, the regulatory mechanisms underlying this dynamic transcriptome remain poorly understood.

Recent studies have challenged the traditional view that repetitive sequences are merely nonfunctional remnants of genomic evolution, revealing that some transposable elements play critical roles in embryogenesis^4–13^. For instance, Mouse Endogenous Retrovirus with Leucine tRNA primer (MERVL) serves as an essential RNA that helps shape the early embryonic transcriptome, particularly during the transition from totipotency to pluripotency^4,5,12^. The presence of MERVL elements within genic regions can contribute novel transcriptional activities or generate alternative isoforms, orchestrating early developmental processes^1,4,9^. Other repetitive elements, such as the *Obox* and *Dux* family paralogous gene clusters, tightly regulate MERVL transcription, thereby promoting ZGA^7,14–21^. In addition, transient expression of the *Zscan4* gene family has been highlighted as a defining feature of telomere lengthening during early embryogenesis, although its precise function remains unclear^22–29^. Despite these advances, the functional significance of many repetitive elements remains elusive, and further investigation is needed to unravel their roles in early mammalian development.

This study identified RNA export factor in early embryos as a novel, redundant yet critical factor in mouse embryogenesis. *Refee*, an abundantly expressed 2C gene, is a paralog of *Alyref,* a nucleocytoplasmic RNA exporter, and preferentially binds to MERVL RNA, promoting its stability and export. Through interactions with adjacent MERVL elements, *Refee* appears to play a dual role: enhancing major ZGA transcript levels including MERVL RNA, while simultaneously repressing most 2C gene transcripts. These findings suggest that *Refee*, in conjunction with other repetitive elements, plays a key role in the proper execution of ZGA at the post-transcriptional level, ensuring early embryonic progression beyond the 2C stage.

## Results

### 1. Discovery of *Refee* as a redundant 2C-specific gene

Gene redundancy allows for the rapid accumulation of transcripts and safeguards against the loss of essential survival mechanisms. ZGA is a fundamental developmental milestone critical for species survival and continuity, during which a broad array of redundant genetic elements, including repetitive sequences and duplicated genes, are actively transcribed^3^ (Fig. 1a). Considerable efforts have been made to elucidate the function of these elements, such as the mouse-specific transcription factors DUX and OBOX4^14–18,20^. However, strikingly, even near-complete loss of DUX and/or OBOX4 results in only modest alterations to the ZGA transcriptome^16,17,20^. This observation supports the idea that mouse ZGA is “so strongly canalized”^20^ that disruption of any single or even a few regulatory pathways is insufficient to halt the complex, multilayered progression of ZGA. Therefore, dissecting the role of each redundant element—even one at a time—remains essential for understanding the robust regulatory architecture of mammalian genome activation.

**Figure 1.**
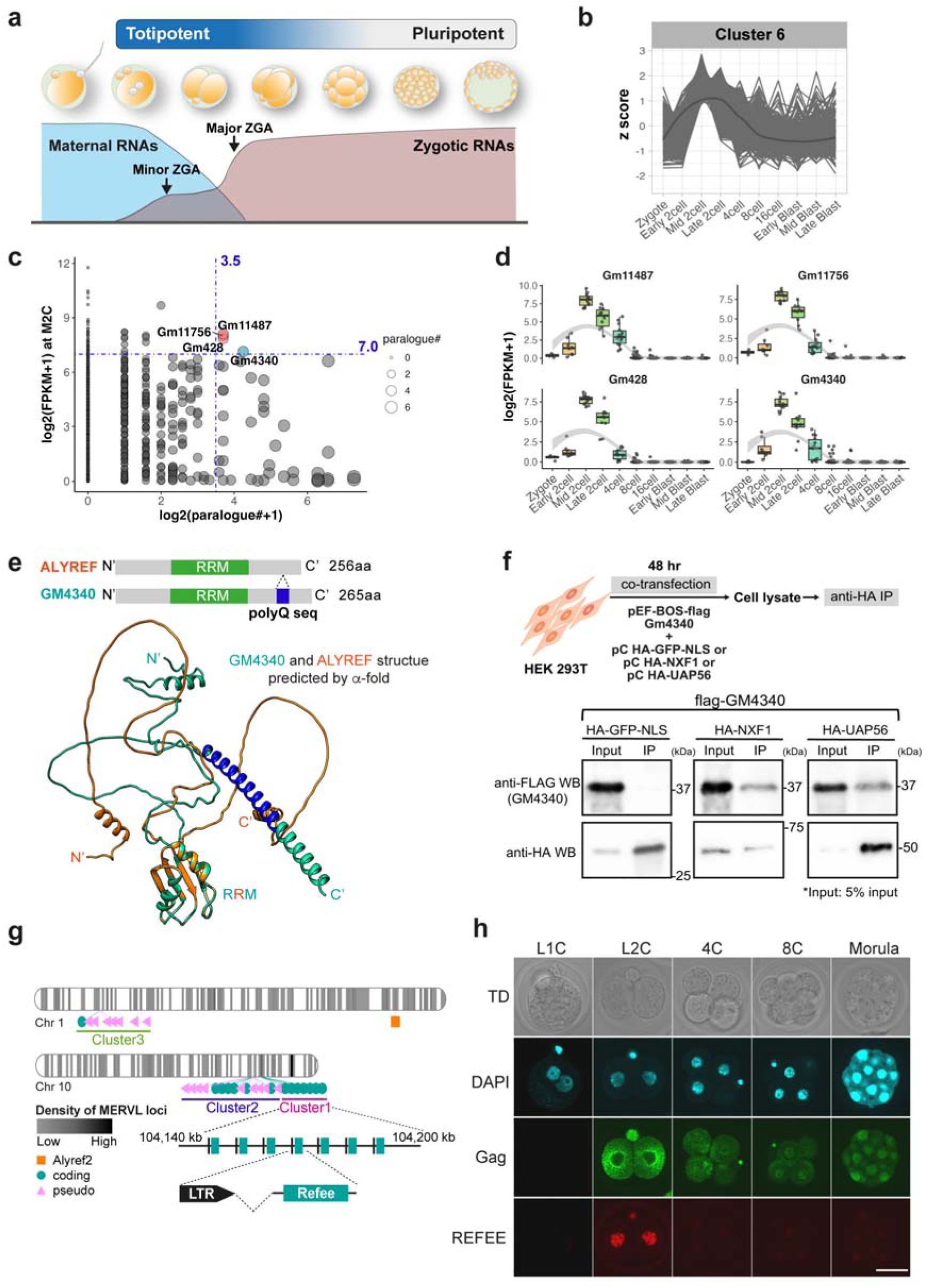
*Refee* is a 2C gene paralog of *Alyref* expanded in mice. **a.** Schematic diagram of mouse ZGA and preimplantation development. After the fertilization, zygotic genome initiates stepwise transcriptional bursts, such as minor and major ZGA occur at the early and late 2-cell (2C) stages, respectively. In contrast, maternal transcripts disappear until the execution of ZGA. **b.** Expression kinetics of cluster 6 genes analyzed using single-cell RNA-seq data by Deng et al.^30^. Blast, Blastocyst **c.** Distribution of 2C genes according to expression levels in mouse M2C embryos and the number of paralogues. FPKM, Fragments Per Kilobase of exon per Million mapped fragments **d.** Expression patterns of the top four highly redundant 2C genes during preimplantation development. **e.** Three-dimensional structures of GM4340 and ALYREF. The upper panel shows the domain architecture of each amino acid sequence. GM4340 uniquely features a polyglutamine sequence at its N-terminal region. The lower panel shows an overlay of structures predicted by AlphaFold and retrieved from the Uniprot database (AF-B9EJS2-F1 for GM4340 and AF-O08583-F1 for ALYREF). The beta-barrel region, corresponding to the RNA recognition motif (RRM), displays a remarkably high similarity. **f.** Interaction between REFEE and RNA export factors. REFEE was detected in immunoprecipitants of UAP56 and NXF1, but not in the nuclear GFP (negative control). **g.** Genomic loci of *Refee* on mouse chromosomes 1 and 10. The chromosomes are colored according to the density of MERVL loci. *Alyref2*, a possible intermediate in the diversification from *Alyref* to *Refee*, is also located on chromosome 1. **h.** Confocal images of embryos from the zygote to morula stages, immunostained for REFEE. Scale bar: 20µm

In this study, we first aimed to identify multicopy 2C genes based on the number of gene paralogues in the mouse genome. To this end, we extracted genes transiently expressed at the mid-2C (M2C) stage, shortly before major ZGA, using publicly available single-cell RNA-seq data from mouse preimplantation embryos^30^ (**Fig. 1b, Fig. S1a**). We then compared transcript abundances (FPKM) of these 2C genes with their numbers of paralogues retrieved from the Ensembl database^31^ (**Fig. 1c**, **Table S1**). Under specific criteria (i.e., > 10 paralogues and log 2 FPKM > 5), four redundant genes were identified: *Gm11487*, *Gm11756*, *Gm428*, and *Gm4340* (**Fig. 1c and d**). The genes are predicted to encode proteins but remain functionally uncharacterized.

Three of them, *Gm11487*, *Gm11756*, and *Gm428*, belong to the *Myb/SANT DNA binding domain containing 5* (*Msantd5*) family. This gene family, specific to certain placental mammals, appears to have expanded in mice, as suggested by a gene family tree inferred using CAFE^32^ (**Fig. S2**). The remaining gene, *Gm4340* (also annotated as *Alyreffm10*), is a paralogue of *Alyref* (*Aly/ RNA Export Factor*), which is broadly conserved among jawed vertebrates. *Alyref* plays a key role in mRNA splicing and export ^33–37^and is essential for embryonic development, as shown by embryonic lethality in homozygous knockout (KO) mice^38^. Although its biological role remains unclear, *Gm4340* has frequently been used as a marker for 2-cell-like embryonic stem cells (2CLCs) in previous studies^7,39^. Its predicted protein product shares 66.6% sequence similarity with ALYREF and contains an N-terminal polyglutamine tract. AlphaFold predictions^40^ revealed high structural similarity in the RNA recognition motifs (RRM) of GM4340 and ALYREF, which are essential for binding exon junction complexes (EJCs)^36^ (**Fig. 1e, Fig. S1b**). Immunoprecipitation analysis showed that GM4340 interacts with known ALYREF-associated mRNA export factors, such as NXF1 and UAP56 (**Fig. 1f**). Based on these data, we hypothesized that this protein functions in RNA export and named this group of gene paralogues *RNA Export Factor interactive in Early Embryos* (*Refee*).

Although 18 *Refee* paralogues are currently annotated in the Ensembl database, manual curation using RetroGenes V6 identified over 30 *Refee* loci on mouse chromosomes 1 and 10, including 17 pseudogenes, which can be classified into three clusters based on genomic loci and RNA sequence homology (**Fig. 1g, Fig. S1c, Table S2**). Two clusters of paralogues were found on chromosome 10 (clusters 1 and 2). Notably, cluster 1 includes copies with upstream sequences highly homologous to the MERVL promoter (MT2_Mm). Phylogenetic analysis of *Alyref* and its paralogues indicates that *Refee* and *Alyref2* diverged from *Alyref* within the subgenus *Mus* (**Fig. S1d**), with *Refee* showing particularly high duplication in *Mus musculus*.

To understand how *Refee* may functionally differ from its paralogues, we next compared their expression patterns during preimplantation development (**Fig. S3a**). *Alyref* expression was initiated at the M2C stage and remained constitutively expressed thereafter, a pattern typical of major ZGA genes^41^. In contrast, *Alyref2* expression gradually decreased throughout development and did not follow the transient expression pattern of *Refee*. To investigate protein expression of REFEE, we performed immunostaining during early embryogenesis using an anti-REFEE antibody generated in this study. REFEE protein was transiently expressed at the 2C stage and localized to the nucleus (**Fig. 1h**). ALYREF was also localized to the nucleus but showed significant expression starting at the 4-cell (4C) stage (**Fig. S3b**).

Collectively, these results indicate that *Refee* encodes a nuclear protein with a distinct temporal expression pattern, potentially acting in totipotency regulation through RNA and EJC interactions.

### 2. REFEE preferentially binds MERVL-derived RNAs

The abobservations strongly suggest that REFEE is an RNA-binding protein capable of regulating the metabolism of its targeted RNAs. To identify REFEE-associated RNAs, we performed individual nucleotide resolution cross-linked immunoprecipitation followed by sequencing (iCLIP-seq)^42^. For this, we used 2CLCs generated by doxycycline (Dox)-induced DUX expression in an embryonic stem cell (ESC) line, TRE::Dux^5,20^ (**Fig. 2a**). REFEE expression was confirmed in both TRE::Dux and spontaneously converted 2CLCs (**Fig. S4a and b**). iCLIP analysis revealed strong interactions between REFEE and transcripts in 2CLC (**Fig. 2a and S4c**). cDNA libraries were generated from the immunoprecipitated complexes and subjected to sequencing. This analysis identified 68,746 binding peaks, nearly half of which (49.2%) mapped to transposable elements (TEs) (**Fig. 2b, S4d, Table S3**). Among the TE-derived peaks, REFEE showed the highest enrichment for MERVL elements, with approximately 10% of all peaks mapping to MERVL-int (internal sequences, ranked #1) and MT2_Mm (LTR promoter sequences, ranked #3) (**Fig. 2c**). These MERVL-associated peaks covered 55.3% of all annotated MERVL loci (**Fig. 2d**), demonstrating REFEE’s preferential binding to MERVL RNAs.

**Figure 2.**
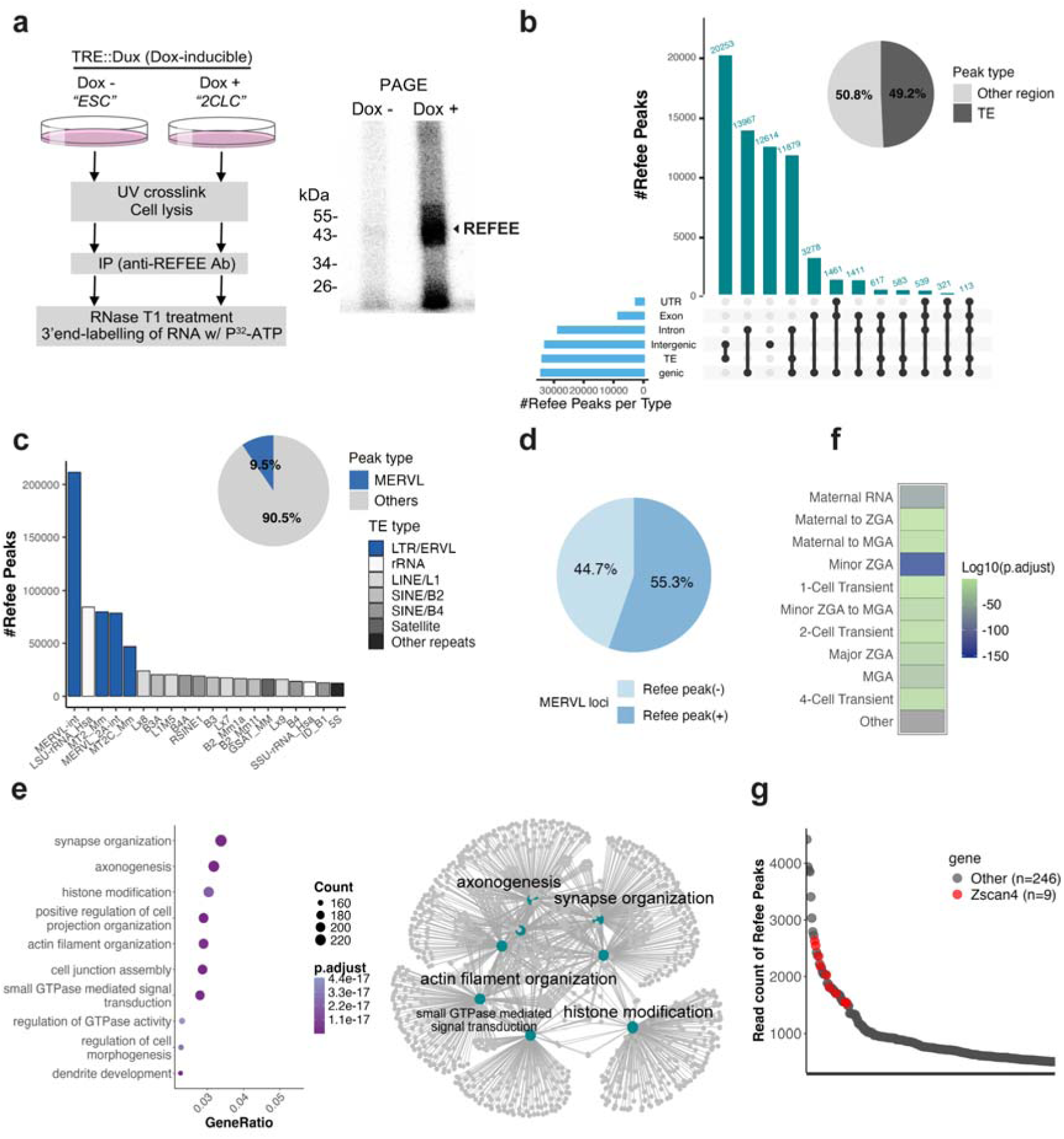
*Refee* is an RNA-binding protein that preferentially targets MERVL RNAs and totipotent embryonic transcripts. **a.** The left panel illustrates the iCLIP scheme. The right panel shows a radiograph of immunoprecipitated products after polyacrylamide gel electrophoresis. A strong radioactive signal at REFEE’s molecular weight was observed in Dox-induced 2CLC (Dox+), while a little signal was detected in ESC (Dox−). cDNA libraries for iCLIP-seq (n = 2) were prepared from the band around 45 kDa. IP, immunoprecipitation **b.** Content of the REFEE binding sites (peaks) identified by the iCLIP-seq. The upset plot shows the overlap between REFEE-bound peaks and genomic regions (n = 68,746, p.adjust < 0.05). The pie chart represents the overall proportion of TE-associated peaks. TE, transposable elements. **c.** TEs targeted by REFEE. The bar plot shows the top 20 TE targets of REFEE. The x-axis represents the total read counts of peaks mapped to each TEs, reflecting REFEE’s binding preference. Blue bars indicate TEs classified as mouse endogenous retrovirus L (MERVL). The pie chart shows the proportion of MERVL peaks among all identified peaks. **d.** Pie chart showing the proportion of each MERVL locus (MERVL-int, MT2_Mm, n=4,075) covered by REFEE-bound peaks. **e.** Enrichment of the REFEE peaks in early embryonic transcripts from the DBTMEE database. Adjusted p-values (p.adjust) were calculated by the hypergeometric test. Non-significant enrichments (p.adjust = 1) are shown in grey. **f.** GO terms enriched in REFEE target genes. The left panel shows the top 10 GO terms on gene coverage within each term (GeneRatio). The right panel visualizes the relationships among the top 8 GO terms according to gene overlap. **g.** Dot plot showing the top 136 REFEE targets ranked by total read counts per peak. *Zscan4* family genes are highlighted in red.

In addition to TEs, REFEE also bound to genic transcripts, including intronic regions (**Fig. 2b and S4e**). Gene Ontology analysis of REFEE-bound genes (n = 1,673) revealed enrichment for functions such as synapse organization and histone modification (**Fig. 2e**). Notably, many of these transcripts overlapped with those enriched in mouse totipotent-stage embryos, as documented in the DBTMEE database^41^ (**Fig. 2f**). Among them, the most prominent targets were mRNAs from the *Zscan4* gene family, a set of multicopy 2C-specific genes (**Fig. 2g**).

Taken together, these results suggest that REFEE may orchestrate key molecular events during early embryonic development by directly interacting with the totipotency-associated RNAs.

### 3. REFEE regulates MERVL RNA expression post-transcriptionally

The iCLIP-seq analysis revealed that REFEE predominantly interacts with MERVL and *Zscan4* RNAs. To investigate the functional role of *Refee* in ESCs, we generated *Refee* KO ESCs by introducing mutations in the *Refee* cluster 1 locus using CRISPR/Cas9-mediated gene targeting (**Fig. 3a**). Since cluster 1 accounts for approximately 96% of total *Refee* transcripts in ESCs, as revealed by absolute quantification of each cluster’s transcripts using RT-qPCR, we focused on editing this cluster (**Fig. S5a–b**). We isolated three mutant clones: clone #14, which harbors a deletion of the entire cluster 1 locus, and clones #11 and #34, each containing a single-nucleotide deletion predicted to cause a frameshift mutation at histidine 81 (**Fig. S5c–e**). These mutations resulted in a marked reduction of *Refee* expression at both mRNA and protein levels (**Fig. 3b, S5f–g**). Importantly, protein and RNA expression levels of *Alyref* were not significantly altered, confirming the specificity of the gene targeting (**Fig. S5h, S6a-b**).

**Figure 3.**
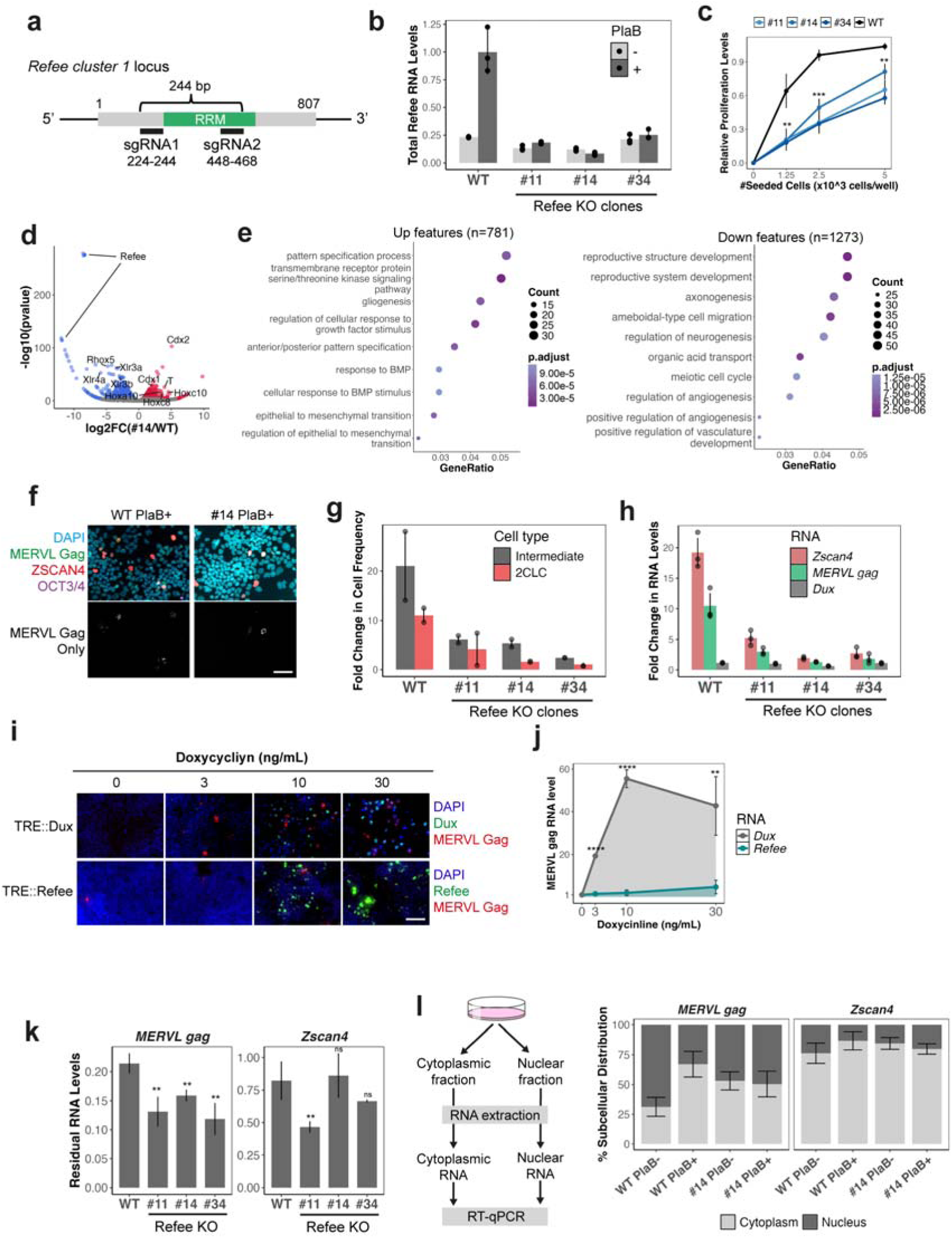
REFEE promotes the conversion of ESCs into 2-cell-like cells. **a.** Schematic showing the position of small guide RNAs (sgRNAs) for the CRISPR/Cas9 gene targeting. **b.** *Refee* RNA expression levels in each ESC clone (#11, 14, and 34) with or without Pladienolide B (PlaB), a compound that induces a 2CLC-like state (n = 3). **c.** Cell proliferation of ESC clones assessed by colorimetric assay 2 days after seeding into 96-well plates (n = 3). **d.** Volcano plot showing differentially expressed genes in clone #14. Blue, downregulated genes; red, upregulated genes. **e.** Top enriched GO terms among upregulated (the left panel) and downregulated genes, excluding *Refee* (the right panel). **f.** Immunofluorescence of OCT3/4, ZSCAN4, and MERVL Gag in wild-type (WT) and clone #14 ESCs after 3 days of PlaB treatment. The upper panel shows merged images of the markers with DAPI. The lower panel displays the MERVL Gag signal alone. Scale bar: 30µm **g.** Efficiency of 2CLC and intermediate states, evaluated by the frequency of marker-positive cells (Fig. S5c) (n = 2). **h.** Fold change in *MERVL gag*, *Zscan4*, and *Dux* RNA levels by the 2CLC induction. **i, j.** Dose-dependent induction of DUX or REFEE. Immunofluorescence analysis after Dox-induced expression of *Dux* or *Refee* in a dose-dependent manner (**i**) and relative expression levels of MERVL RNA following Dox-induced expression of *Dux* or *Refee* (**j,** n = 3). TRE::Dux or TRE::Refee ESC lines were used. Scale bar: 30µm. **k.** Stability of MERVL and *Zscan4* RNAs in *Refee* KO ESCs after 2CLC induction. Relative RNA levels remaining at 3 hours (hr) post-Actinomycin D treatment compared to 0 hr were plotted (n = 3). **l.** Cytoplasm and nuclear location of MERVL RNA. The left panel illustrates the experimental design. The right panel shows the percentage of cytoplasmic and nuclear distribution, calculated based on the relative RNA abundance in each fraction (n = 2). Statistical significance was evaluated using the student’s t-test in **b**-**c**, **h**, and **j-k** (**p<0.05, ***p<0.01, ns; non-significant). Bars on the plots represent mean ± SD in **b**, **c**, **g**, **h**, and **j-l**.

All three KO clones exhibited reduced proliferation compared to wild-type ESCs, with their relative growth ranging from 0.25 to 0.76 (**Fig. 3c**). Transcriptome analysis of clone #14 showed altered expression patterns of several developmental regulator genes, including *Cdx* family members, *Hox* genes, and *T* (*Brachyury*) (**Fig. 3d, 3e, S6a-b, Table S4**), suggesting broader perturbation of the developmental gene network. These transcriptional changes are indicative of a potential disruption in cell identity regulation, which could affect the dynamic transitions between pluripotent and totipotent-like states.

Given the role of developmental gene networks in cell fate determination, we next examined whether loss of *Refee* affects 2CLC conversion efficiency. Immunostaining for ZSCAN4 and MERVL Gag allowed quantification of 2CLC (ZSCAN4+/MERVL+) and intermediate (ZSCAN4+/MERVL–) populations^43^ (**Fig. 3f**). While the proportion of 2CLC remained unchanged in KO clones, the intermediate population was significantly increased relative to wild-type ESCs. Upon 3-day treatment with Pladienolide B (PlaB), an inducer of totipotent blastomere-like cells (TBLC)—a specific 2CLC state—^44^, 2CLC frequencies decreased in mutant clones, whereas the intermediate population remained largely unaffected (**Fig. S6c**). The 2CLC conversion efficiency, calculated as the ratio of 2CLC frequency in PlaB-treated versus untreated cells, was consistently reduced in all KO clones (**Fig. 3g**). Consistently, MERVL *gag* and *Zscan4* transcript levels showed a diminished response to PlaB-induced 2CLC conversion, despite stable expression of *Dux*, a known transcriptional activator of these genes ^14,15^ (**Fig. 3h**).

To further explore *Refee*’s regulatory role, we performed ectopic expression experiments. While ectopic DUX expression strongly induced MERVL, ectopic expression of REFEE alone did not alter MERVL transcript levels (**Fig. 3i, j**), suggesting a post-transcriptional mechanism of regulation. Indeed, truncation of REFEE markedly decreased the stability of MERVL RNA (**Fig. 3k**) and impaired its nucleocytoplasmic export (**Fig. 3l**). In contrast, *Zscan4* mRNA showed minimal changes in both stability and export upon REFEE loss, indicating that distinct regulatory mechanisms govern these two RNA targets.

### 4. REFEE stabilizes MERVL RNA and supports early embryonic development

To investigate whether *Refee* plays a functional role in early embryos, we first examined the subcellular localization of REFEE and MERVL RNA. Consistent with the findings in ESCs, immunostaining combined with smRNA-FISH revealed prominent colocalization of REFEE and MERVL RNA in early- to mid-2C stage embryos (**Fig. 4a**), indicating that REFEE targets MERVL RNA in vivo.

**Figure 4.**
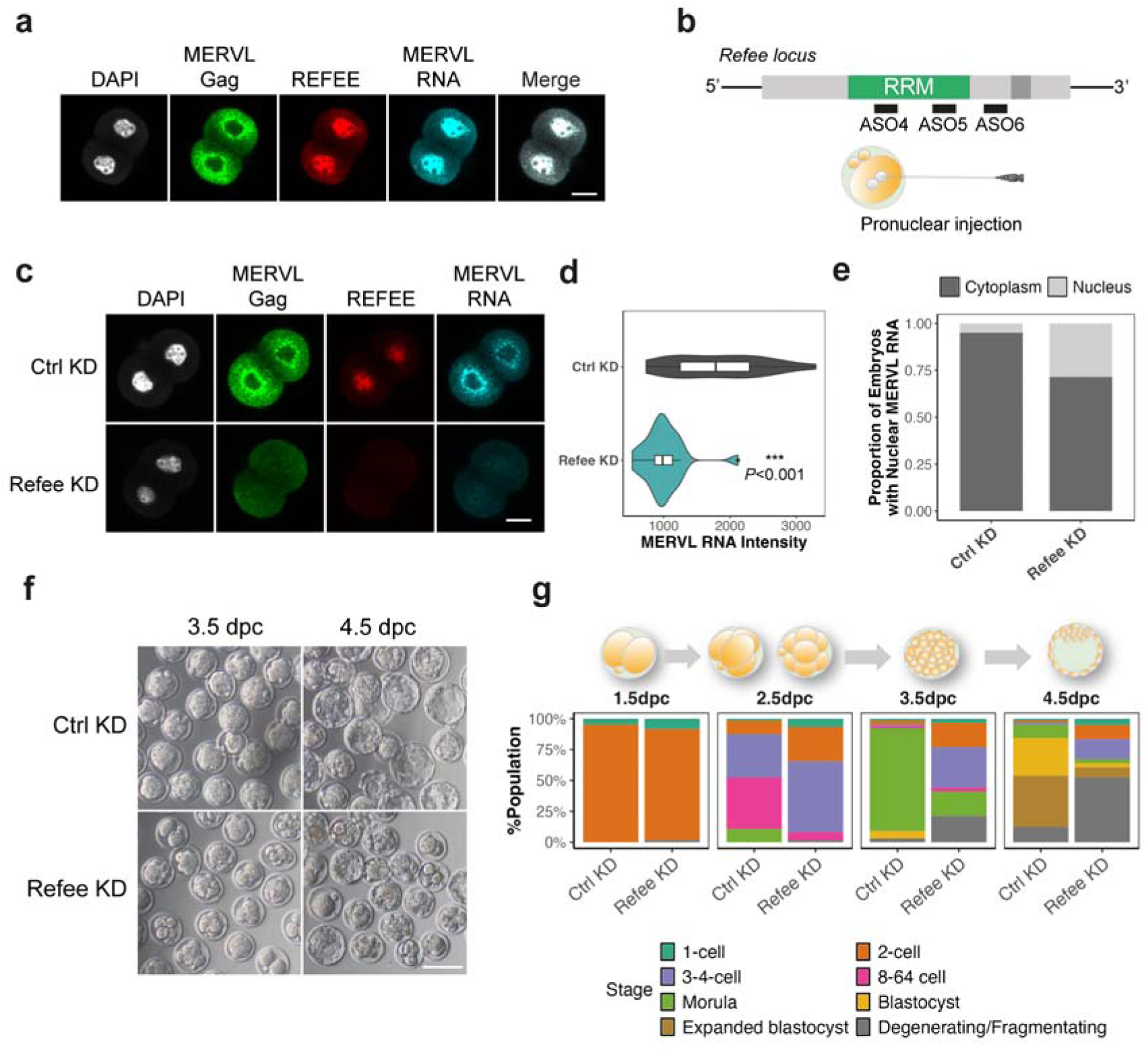
*Refee* knockdown causes developmental arrest in early embryos. **a.** Nuclear colocalization of REFEE and MERVL RNA in the M2C embryos. MERVL Gag and REFEE proteins were immunostained, followed by single-molecule RNA FISH (smRNA-FISH) for MERVL RNA. The merged image shows an overlay of DAPI, REFEE, and MERVL RNA signals. TD, transmitted light image. Scale bar: 20µm **b.** Experimental design for *Refee* knockdown (KD) in embryos. **c.** Knockdown efficiency confirmed by immunostaining combined with smRNA-FISH. Representative images of KD L2C embryos. Scale bar: 20µm **d.** Signal intensity of MERVL RNA in KD L2C embryos from **c** (n = 25 for control, n = 21 for *Refee* KD). Bars represent median values ± SD. Statistical significance was calculated by the Wilcoxon rank sum test. **e.** Proportion of embryos with nuclear MERVL RNA, calculated based on the data shown in **c**. **f.** Representative images of KD embryos at 3.5 and 4.5 dpc. Scale bar: 50µm **g.** Developmental status of KD embryos. Proportions of developmental stages in 65 control embryos (Ctrl) and 61 *Refee* KD embryos from two respective experiments.

To assess the functional relevance of *Refee* during preimplantation development, we performed knockdown (KD) experiments by injecting a mixture of three antisense oligonucleotides (ASOs) into the paternal pronucleus of mouse zygotes (**Fig. 4b**). Although two of the ASOs (ASOs 4 and 6) were capable of targeting *Alyref* transcripts, we confirmed that their administration led to a substantial reduction of REFEE protein levels in both embryos and ESCs, while ALYREF protein levels remained largely unaffected (**Fig. 4c, Fig. S7, S8a-b**). Single molecular RNA-FISH of MERVL RNA combined with MERVL Gag immunostaining revealed that depletion of REFEE markedly impaired MERVL expression in late-2C (L2C) embryos (**Fig. 4c and d**). In addition, the MERVL RNAs exhibited defective nucleocytoplasmic export, which normally occurs during the mid- to late-2C stage^5^ (**Fig. 4e**). These results are consistent with our ESC-based findings, indicating that REFEE stabilizes MERVL RNA and promotes its cytoplasmic localization in early embryos.

The developmental status of *Refee* KD embryos was monitored over 4.5 days post coitum (dpc), revealing a pronounced arrest at the 2C to 4C stage (**Fig. 4f, g**). The developmental delay was evident from 3.5 dpc onward, with the appearance of arrested embryos. In addition, degeneration became apparent from 4.5 dpc, with the arrested embryos showing extensive fragmentation. In contrast, most control KD embryos continued to develop and reached the blastocyst stage, marking the completion of preimplantation development. The severity and early onset of developmental failure in *Refee* KD embryos suggest a critical role for *Refee* in facilitating embryonic progression beyond the 2C stage.

### 5. *Refee* is required for mouse zygotic genome activation

To investigate the molecular basis of the developmental arrest observed in *Refee* KD embryos, we analyzed the transcriptomes of embryos at the M2C, L2C, and 4C stages using RNA-seq (**Fig. S8a**). Expression analysis of transposable elements revealed a marked reduction in MERVL RNAs, such as MERVL-int and MT2_Mm, upon *Refee* KD, consistent with the smFISH results shown in Figs. 4b and 4c (**Fig 5a**). Subsequent gene expression profiling identified 2,939 differentially expressed genes (DEGs) in *Refee* KD embryos across the M2C to 4C stages (**Fig. 5b, S8c, Table S5**). Further examination of expression patterns revealed that the majority of DEGs were specifically dysregulated at the 4C stage. Notably, the upregulated DEGs were significantly enriched for maternal transcripts, while the downregulated genes included a large number of transiently expressed 4C genes, indicative of developmental delay in the *Refee* KD embryos (**Fig. S8d**). This interpretation was further supported by the reduced expression of pluripotency-related genes such as *Nanog*, *Myc*, and *Esrrb* (**Fig. S8e**).

**Figure 5.**
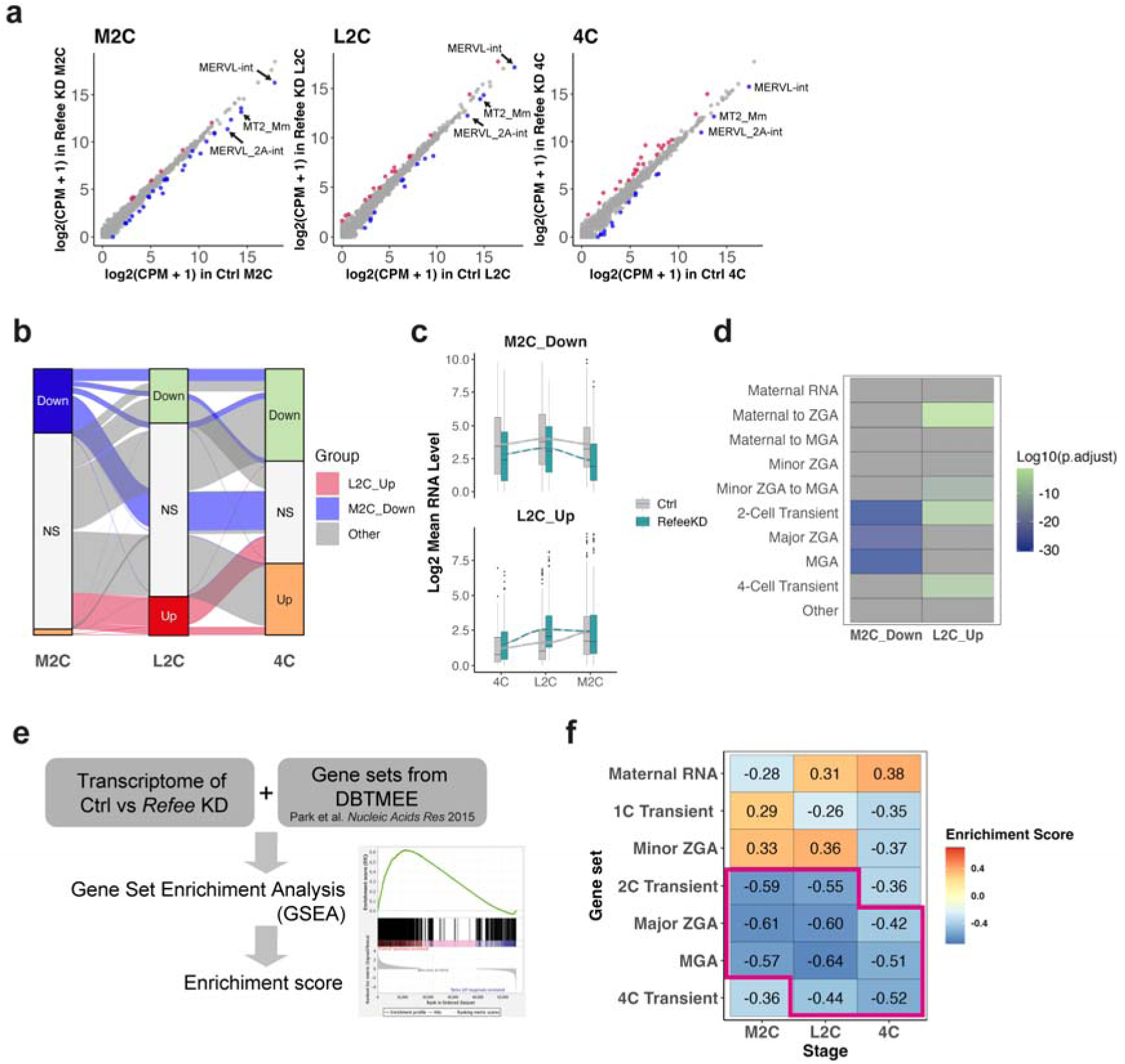
*Refee* is essential for zygotic genome activation in mice. **a.** Scatter plot showing pre-gene normalized read count on TEs in the indicated stage embryos and highlighting differentially expressed TEs after normalization (upregulated in red, downregulated in blue). **b.** Stage-specific expression transitions of the identified differentially expressed genes (n = 2,939). Genes categorized as M2C_Down are highlighted in blue, and L2C_Up genes in red. **c.** Expression patterns of M2C_Down and L2C_Up genes in *Refee* KD embryos. The curves represent smoothed expression trajectories fitted using loess regression. **d.** Enrichment analysis of each gene group, tested by the hypergeometric test. **e, f.** Gene set enrichment analysis (GSEA) using DBTMEE gene sets (**e**) and heat map visualization of enrichment scores (**f**). Negative scores represent downregulated features in *Refee* KD embryos, while positive scores indicate upregulated features. Statistically significant enrichments (q-value < 10L³) are boxed in magenta.

Among the differentially expressed genes, we chose to focus on two distinct subsets with notable expression patterns (**Fig. 5b**). The first subset, M2C_Down (n=675, excluding *Refee*), represents a major population of genes that were significantly downregulated at the M2C stage. The second, more unique subset, L2C_Up (n=427), comprises genes identified as upregulated in DEG analysis but that, based on kinetic analysis, failed to downregulate at the M2C stage and thus persisted at the L2C stage (**Fig. 5c**). These distinct gene subsets, M2C_Down and L2C_Up, were significantly enriched in gene categories such as 2C-transient, major ZGA, and mid-preimplantation genome activation (MGA, which typically occurs at the 4C stage) for M2C_Down, and 2C- and 4C-transient genes for L2C_Up (**Fig. 5d**). These findings indicate that M2C_Down genes play a major role in the failure of major ZGA, and that *Refee* KD embryos also exhibit delayed decay and/or silencing of 2C- and 4C-transient transcripts. Indeed, transcriptome-wide gene set enrichment analysis (GSEA) at each developmental stage consistently revealed significant impairments in major ZGA and MGA gene programs (**Fig. 5e, f, S9**), reinforcing the notion that *Refee* plays a pivotal role in orchestrating ZGA, and this may lead to the 2C to 4C arrest.

### 6. REFEE iCLIP targets coincide with transcriptomic changes and overlap with MERVL-regulated genes

To investigate how REFEE depletion leads to both downregulation and upregulation of specific gene sets, we turned our attention to MERVL, given its prominence in the iCLIP-seq data and the *Refee* KD phenotype. Previous study has shown that depletion of MERVL can lead to the upregulation of nearby genes^5^, suggesting potential cis-regulatory roles. We therefore examined the genomic proximity between MERVL elements, including internal sequences (MERVL-int) or LTRs (MT2_Mm), and the REFEE-regulated gene sets. M2C_Down genes exhibited the shortest median distance to the nearest MERVL elements among the four groups (**Fig. 6a**) and were significantly more likely to harbor MERVL insertions within their gene bodies (**Fig. 6b)**. This included various MERVL-driven chimeric genes, such as *Aqr*, *Piwil2*, and *Pemt* (**Fig. 6c**), suggesting that loss REFEE destabilizes these transcripts. In contrast, L2C_Up genes also tended to be relatively close to MERVL elements, often located in intergenic regions within gene clusters such as *Usp17l* and *Zscan4* (**Fig. 6d**), raising the possibility that loss of REFEE derepress nearby gene expression via MERVL RNA decay.

**Figure 6.**
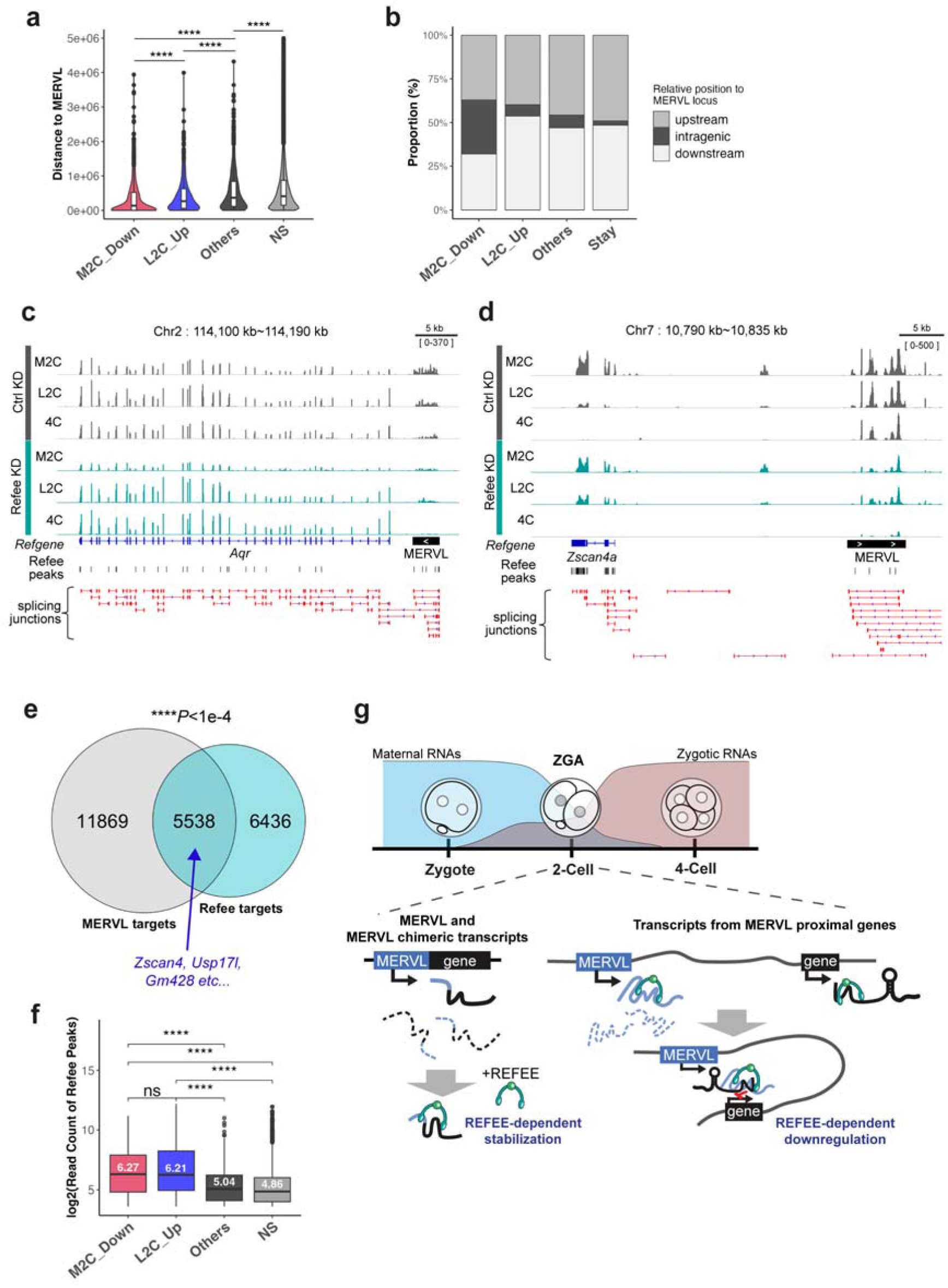
*Refee* exerts its function through the regulation of MERVL–derived RNAs. **a, b.** Positional relationship between genes and MERVL loci. (**a**) Distance to the nearest MERVL locus for genes in the M2C_Down and L2C_Up groups. (**b**) Positional distribution by type. **c, d**. Genome browser snapshots of a representative M2C_Down gene, *Aqr* (**c**), and L2C_Up genes, *Zscan4a* (**d**), together with their adjacent MERVL elements. **e.** Overlap between REFEE binding targets and gene silencing targets of MERVL RNA. Statical significance was tested by the hypergeometric test. **f.** Bar plot showing the log_2_(read count of REFEE peaks on each gene), grouped by RNA expression change categories: “M2C_Down”, “L2C_Up”, “Others” (differentially expressed genes not included in M2C_Down or L2C_Up), and “Stay” (genes without significant RNA expression changes) (from Fig. 5b). The numbers in each box represent the median value. Statistical significance was assessed using the Wilcoxon rank sum test (****, p < 0.001; ns, not significant). **g.** Conceptual model illustrating REFEE’s post-transcriptional regulation mediated via MERVL transcripts. The RNA-binding protein REFEE performs two distinct, seemingly contradictory roles that together orchestrate the early embryonic transcriptome via MERVL transcripts. First, REFEE stabilizes and facilitates the nuclear export of MERVL RNA and its chimeric transcripts, promoting the expression of MERVL-driven genes (the left panel). Second, REFEE acts as a repressor; its binding to a MERVL transcript can lead to the silencing of a nearby 2C gene transcript (the right panel). This cis-regulatory mechanism may also result in a loss of chromatin accessibility in the region. Thus, REFEE’s dual function— both promoting MERVL transcription and simultaneously repressing nearby 2C genes—may cooperatively fine-tune the gene expression profile necessary for early embryonic development.

We next sought to understand the effects of MERVL RNA loss by comparing the gene expression changes in *Refee* KD embryos with those observed in our previous MERVL KD study^5^. Both conditions led to significant degeneration at 4.5 dpc; however, *Refee* KD embryos appear arrested at a significantly earlier stage than those observed with MERVL KD. Despite this difference in developmental timing, the comparison of their differentially expressed genes showed overlapping but distinct sets of genes (**Fig. S10a, b and Table S7**). Among 692 M2C_Down genes in *Refee* KD embryos, 85 (12.3%) were also downregulated in MERVL KD embryos, while 607 (87.8%) were uniquely affected by *Refee* KD, suggesting both MERVL-dependent and -independent functions of REFEE. In addition, we found a significant overlap between REFEE binding targets and genes associated with downregulated ATAC-seq peaks in MERVL KD embryos (**Fig. 6e and Table S7**), indicating a model in which REFEE contributes to MERVL RNA-mediated regulation of chromatin accessibility. Finally, we observed that REFEE binding targets were significantly enriched among the M2C_Down and L2C_Up gene groups (adjusted p-values = 1.34e–45 and 2.05e–12, respectively), and these genes also exhibited significantly higher REFEE-iCLIP signal intensities compared to other groups, indicating that REFEE preferentially binds transcripts from both groups (**Fig. 6f**).

Altogether, these findings support a model in which REFEE controls zygotic transcript levels through two distinct modes: by stabilizing MERVL RNAs to maintain expression of adjacent genes, and by repressing gene expression near derepressed MERVL loci, possibly through the loss of MERVL-derived regulatory (**Fig. 6g**).

### 7. Impact of *Refee* on proteome change in early embryos

To assess whether *Refee* depletion affects the proteome, we next performed mass spectrometry on KD L2C embryos, the developmental stage when major ZGA begins and embryonic arrest is first observed upon *Refee* loss. Using a tandem mass tag (TMT)– based quantitative approach, we quantified 4,228 proteins, corresponding to approximately 30% of the mouse reference proteome. The majority of these identified proteins were maternally deposited or encoded by minor ZGA genes —those expressed before L2C stage (**Fig. 7a**). Statistical analysis identified 11 proteins that were significantly upregulated and 66 proteins that were downregulated by *Refee* depletion, including MERVL Gag, which was among the most strongly reduced proteins (**Fig. 7b and Table S6**). Among the upregulated proteins, three were involved in ubiquitination, although this enrichment was not statistically significant. In contrast, downregulated proteins were significantly enriched in nuclear proteins (n = 41), particularly those involved in transcription initiation from the RNA polymerase II promoter (FDR = 0.0121, Enrichment strength = 1.52) and RNA metabolic processes (FDR = 0.0141, Enrichment strength = 1.24) (**Fig. 7c**). Despite the limited number of differentially expressed proteins, they were significantly enriched among maternally expressed and major ZGA genes (**Fig. 7d**).

**Figure 7.**
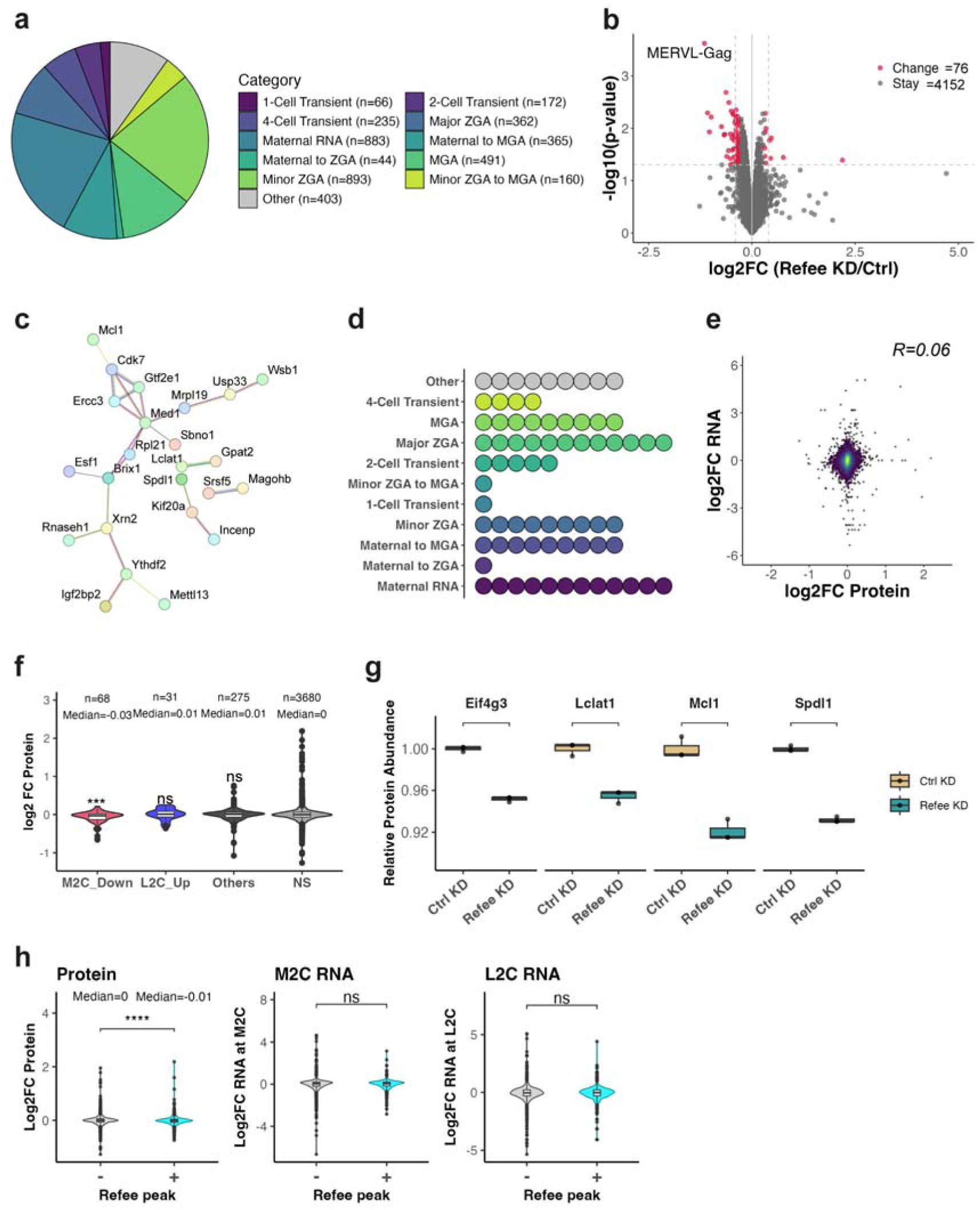
*Refee* regulates target genes at both RNA and protein levels. **a.** Attribution of the quantified proteins based on embryonic gene categories. The protein quantified in both *Refee* KD and control L2C embryos using tandem mass tag (TMT) method were aligned to DBTMEE database^41^. **b.** Volcano plot of protein expression changes between *Refee* KD and control L2C embryos, based on the proteomics data. **c, d**. Functional protein association networks (**c**) and DBTMEE category (**d**) of differentially expressed proteins identified in **b**. One dot represents one differentially expressed protein in **d**. **e.** Scatter plot showing the weak correlation between RNA and protein expression changes in *Refee* KD L2C embryos (Pearson’s r (R) =0.06). **f.** Violin plot showing the log_2_ FC of protein level between *Refee* and control KD embryos grouped by RNA expression change categories. Asterisks represent statistical difference compared with the “Stay” group, assessed by the Wilcoxon rank sum test (****, p < 0.001; ns, not significant). **g.** Expression level of representative M2C_Down proteins in the KD embryos. **h.** Violin plots depicting expression changes grouped by REFEE interaction (REFEE iCLIP peak +/-). Left, protein level changes; middle, RNA level changes in KD M2C embryos; right, RNA level changes in KD L2C embryos. Statical significance between the groups was calculated by the Wilcoxon rank sum test in **g-h** (****, p < 0.001; ***, p < 0.01; ns, not significant).

It is well established that transcriptome and proteome profiles are poorly correlated in early embryos prior to the completion of ZGA^45–47^. Consistent with this, the overall correlation between transcript and protein log fold changes (logFC) in *Refee* KD versus control embryos was extremely weak (**Fig. 7e**). To further dissect this relationship, we examined proteomic changes within the DEG groups identified in Fig. 5b, revealing significant downregulation in the M2C_Down group, whereas the L2C_Up group showed no substantial change (**Fig. 7f, g**). These findings suggest that the decreased protein abundance of M2C_Down genes may play a causal role in the developmental arrest observed between the 2C and 4C stages, while L2C_Up gene products unlikely drive the arrest phenotype, given that *Refee* KD embryos begin to arrest as early as the 2C stage.

To further explore the connection between REFEE binding and protein expression, we classified genes into REFEE-bound (iCLIP+) and non-bound (iCLIP−) categories. Interestingly, the iCLIP+ group exhibited significantly reduced protein levels in *Refee* KD embryos, whereas their transcript levels remained unaffected at both M2C and L2C stages (**Fig. 7h**). These results indicate that REFEE binding may promote the expression level of a subset of zygotic proteins without changing its transcript level, in addition to the stabilization of MERVL-associated transcripts.

## Discussion

ZGA is a pivotal developmental transition that has been extensively studied from the transcriptional regulation perspective (reviewed in ^3^). While several factors, including m6A writers^48^, splicing regulators^49^, and minor spliceosome components^50^, have been implicated in ZGA, most function through general RNA processing pathways or maternal transcript clearance. In contrast, our study identifies REFEE, a repetitive element-derived, mouse-specific paralogue of ALYREF^33–37^, as a post-transcriptionally regulator of zygotic transcripts —including retrotransposon RNAs— through a developmentally coordinated program of stabilization and export. Loss of REFEE leads to dysregulated transcript levels and impaired ZGA progression, revealing a previously unrecognized layer of regulatory control embedded within the repetitive genome.

Specifically, we uncovered REFEE’s dual, seemingly paradoxical roles that collectively orchestrate the early embryonic transcriptome: (1) stabilizing and facilitating the export of MERVL RNA, and (2) repressing nearby 2C gene transcripts. These spatially coordinated functions exploit the genomic proximity between MERVL loci and adjacent 2C genes, representing a previously unrecognized post-transcriptional regulatory mechanism mediated by a repetitive element. However, this repression is embryo-specific and is not recapitulated in 2C-like ESCs, highlighting the developmental context-dependency of this mechanism. Thus, the specific role and significance of REFEE in the maintenance of ESCs warrant further investigation.

We observed that loss of REFEE reduces MERVL transcripts in embryos and ESCs, supporting REFEE’s involvement in stabilizing MERVL RNA and promoting its cytoplasmic export. This finding contrasts with the phenotypes of exosome-deficient ESCs, where MERVL RNA accumulates due to impaired degradation, leading to 2CLC states^51,52^. This observation, in conjunction with our data, suggests that REFEE stabilizes MERVL transcripts by antagonizing nuclear exosome-mediated degradation, potentially through competition as similarly reported for ALYREF^53^. Such regulation adds new insight into how early embryonic cells manage retrotransposon RNA surveillance post-transcriptionally.

Given REFEE’s homology to the RNA export adaptor ALYREF and its interactions with RNA export machinery, its role in nuclear export is plausible yet temporally restricted. We show that in early- to mid-2C stage embryos, REFEE and MERVL RNAs colocalize within the nucleus, with MERVL transcripts becoming cytoplasmic only at the late 2C stage. This developmental timing coincides with the absence of canonical RNA export factors such as NXF1 and UAP56 (Fyttd1), which are not expressed until the 4C stage, suggesting that REFEE acts as an early-acting, stage-specific regulator of RNA fate—balancing stabilization and export during a window when the canonical export machinery is not yet functional.

Beyond its role in RNA stabilization and export, REFEE’s regulatory system appears to be more complicated. Based on the mass spectrometry results, despite a low number of statistically significant differentially expressed proteins, we observed a global decrease in the protein levels of REFEE-bound RNAs, even though no changes were detected at the RNA level. This finding suggests that REFEE’s regulatory system may not only influence RNA levels but also affect protein levels exclusively. While one such mechanism could be the control of nucleocytoplasmic transport (**Fig. 3l and 4e**), it is plausible that REFEE also harbors a similar unidentified function to its paralog, ALYREF, which has been reported to possess diverse functions, including splicing promotion^33^, as well as RNA export activities ^34–36^ and co-transcriptional regulation of RNA levels^54^. For instance, REFEE may impact on spliceosome in early embryos, which could then also contribute to the translational control without a change in RNA levels, the regulation of ZGA, and/or the developmental progression beyond the 2C stage. Therefore, continued research on REFEE’s functions will be critical for elucidating the mechanisms of post-transcriptional regulation that govern ZGA.

From an evolutionary perspective, REFEE’s mouse-specific emergence and co-evolution with the MERVL retrotransposon highlight how lineage-specific repetitive elements can be repurposed to acquire indispensable regulatory functions during conserved developmental processes. Unlike its paralogue ALYREF, which regulates post-transcriptional events at later stages such as *Nanog* expression during blastocyst formation^38^, REFEE’s expression is tightly coupled to that of MERVL, making it crucial for the earliest ZGA phase. This highlights functional divergence and species-specific innovation.

In summary, our study reveals REFEE as a dual-function post-transcriptional regulator essential for shaping the early embryonic transcriptome through stabilization, export, and repression of repetitive element-derived RNAs. These findings provide a new conceptual framework for understanding post-transcriptional regulation in mammalian ZGA and highlight the evolutionary significance of repetitive element domestication in early development.

## Methods

### Animals

Male and female B6D2F1 and BALB/c mice were purchased from Japan SLC Inc. The mice were fed regular chow and housed in a controlled room under a 14hr / 10hr light / dark cycle at 23°C ± 3°C, with a humidity of 50% ± 10%. All animal experiments were approved by the Animal Care and Use Committee of Keio University (#A2022-143 and A2022-193).

### Cell lines

HEK293T cells were cultured in 293T medium constituted as follows: Dulbecco’s Modified Eagle Medium (DMEM) (#08459-64, Nacalai Tesque, Japan) supplemented with 10% fetal bovine serum, 1 ✕ GlutaMAX (Thermo Fisher Scientific, USA), 1 ✕ non-essential amino acids (Thermo Fisher Scientific, USA), 1 ✕ sodium pyruvate (Thermo Fisher Scientific, USA), 50µM 2-mercaptoethanol (Thermo Fisher Scientific, USA).

All mouse ESCs (derived from the EB3) were cultured in 293T medium supplemented with 1µM CHIR99021 (Cayman Chemical, USA), 1µM PD0325901 (Cayman Chemical, USA), and mouse leukemia inhibitory factor produced in-house. Culture vessels were coated with porcine gelatine (Sigma, USA) for routine maintenance. For subsequent experimental conditions, iMatrix-511 silk (recombinant human laminin-511 E8 fragment, Matrixome, Japan) was directly incorporated into the culture medium at a concentration of 125 ng/cm².

The above cell lines were all cultured at 37 °C with 5% CO2.

### Plasmids

Plasmids used in this study are listed in Table S8. Doxycycline (Dox)-inducible *Refee* expression piggyBac vector, pTRE-mVenus-Refee-Neo, was generated by swapping the 3FLAG-Dux sequences of pPB-TRE-3FLAG-Dux ^55^ into the N’-mVenus-fused Refee sequence using NEBuilder HiFi DNA Assembly Master Mix (NEB).

3FLAG-fused REFEE and ALYREF expression plasmid, pEF-BOS-flag-Refee and pEF-BOS-flag-Alyref, respectively, were generated by inserting the multiple cloning site of pEF-BOS-bsr^56,57^. HA tag-fused nuclear export factor expression plasmid was constructed using the pCAGGS vector established previously^58^.

### Generation of monoclonal antibodies

The anti-REFEE-RRM (#8-4), anti-REFEE-C’ (#9-1), and anti-ZSCAN4B (#18-7) monoclonal antibodies were produced as previously described^20^. Briefly, the target proteins of interest (POIs), either as maltose-binding protein (MBP)-fusions or glutathione S-transferase (GST)-fusions, were produced by expression in *E. coli* (BL21 strain) (see Table S8 for the expression vector information) and purified by affinity chromatography using Glutathione Sepharose 4B (Cytiva) or Amylose Resin (NEB), respectively. For immunization, BALB/c mice were intraperitoneally immunized with the respective purified MBP-fused POIs emulsified in TiterMax Gold adjuvant (TiterMax USA, Inc.). The mice were routinely immunized until sera tested positive for the POIs. Splenocytes were collected a week after the final booster immunization and fused with SP2/0-Ag14 myeloma cells. Fused cells were then cultured in hypoxanthine-aminopterin-thymidine (HAT) medium for 10 days to select hybridomas. Hybridomas were screened using ELISA against the respective purified GST-fused POIs. Specifically, anti-REFEE antibody-producing hybridoma clone #8-4 was subsequently confirmed not to recognize either C’-terminal or N’-terminal recombinant GM4340 protein (both lacking the RRM domain), indicating its specificity for the RRM domain of REFEE. Positive hybridomas were cloned by limiting dilution and expanded to obtain antibody-containing supernatants.

N-terminal region of REFEE recognizing mouse serum (anti-REFEE-N’) was obtained by intraperitoneal immunizations of the purchased KLH-conjugated peptide (NH2-C+TKIQQRRHDRPDSR, 19 ∼ 32 amino acid position of GM4340, Eurofins, Japan).

### Immunoprecipitation

Plasmid transfections into HEK293T cells were performed by the calcium phosphate method as described previously^59^. Briefly, 383 fmol of HA-fused nuclear export factor expression plasmid and 1.5 μg pEF-BOS-flag-Refee or pEF-BOS-flag-Alyref were co-transfected into the cells seeded on a 6-well plate. After 48 hours (hr), the cells were harvested and lysed with IP buffer (20 mM Tris-HCl (pH 7.4), 150 mM NaCl, 0.1% NP-40, and Protease inhibitor cocktail (Nacalai, Japan)) with a sonication (Branson sonifier SFX150, Emerson Electric Co., USA). Dynabeads Protein G (Thermo Fisher Scientific, USA) conjugated with anti-HA antibody (HA-7, Sigma) were incubated for 1 hr at 4°C and then washed four times with IP buffer. The beads were resuspended in 2 ✕ Laemmli SDS sample buffer (18.75% Glycerol, 250 mM Tris-HCl (pH 6.8), 4% Sodium dodecyl sulfate, and 0.2 mg/mL Bromophenol blue), and their supernatant was subsequently analyzed by Western blot.

### Western blot

The cell lysate was mixed with 2 ✕ Laemmli SDS sample buffer, denatured at 95 °C for 5 min, and separated by 10 or 12.5% SDS-PAGE. The proteins were then transferred onto a Protean Nitrocellulose Membrane (0.45 μm pore size, GE Healthcare, USA) via Power Blotter-Semi-dry Transfer System (Thermo Fisher Scientific, USA). The membrane was blocked with 0.1∼0.5% skim milk (Morinaga, Japan) in PBS-T (0.05% Tween 20) for 15 min at room temperature (RT) with gentle rocking following the incubation with specific antibodies. The signal of horseradish peroxidase conjugated to the antibodies was visualized using ECL Western Blotting Detection Reagents (GE Healthcare). The list of antibodies used in this study is provided in Table S8.

### iCLIP-seq

Library construction for iCLIP-seq was performed as previously described by Murano et al.^60^ with several modifications to pull down RNAs interacting with REFEE. Briefly, 2 ✕ 10^7^ Dox-treated TRE::Dux cells were harvested, washed with PBS, and cross-linked by 254 nm UV irradiation at 200 mJ/cm^2^ using a UV Stratalinker 1800 (Stratagene, USA). The cells were then pelleted, snap-frozen in liquid nitrogen, and lysed with high-salt IP buffer (20 mM Tris-HCl (pH 7.4), 500 mM NaCl, 0.1% NP-40, 2 mM DTT, and Protease inhibitor cocktail (Nacalai, Japan)), followed by sonication. The lysate was clarified by centrifugation at 16,500 *g* for 1 min and treated with 1 U/μl of RNaseT1 (Roche, USA) for 15 min at RT.

RNA-protein complex was immunoprecipitated using anti-REFEE-C’ monoclonal antibody and 50μl of Dynabeads Protein G. The beads were then washed three times with IP wash buffer (20mM Tris-HCl (pH 7.4), 750mM NaCl, 0.1% NP-40) and resuspended IP wash buffer supplemented with 1 U/μl of RNaseT1 for a second round of RNA shearing (15 min at RT). The beads were washed three times with IP wash buffer and once with PNK wash buffer (50 mM Tris-HCl (pH 7.5), 50 mM NaCl, and 10 mM MgCl2). Following washes, the RNA-protein complexes on the beads were treated with T4 PNK (New England Biolabs) in 500mM imidazole-HCl (pH 6.0), 100 mM MgCl2, 50 mM DTT, and 1/50 volumes of RNasin (Promega) for 30 min at 37 °C. After the reaction, the beads were washed three times with PNK wash buffer and subjected to 3’ adaptor ligation.

The RNAs on the beads were end-labeled with γ-^32^P-ATP (PerkinElmer, #NEG502Z) using T4 PNK treatment for 30 min at 37 °C, then separated by 4-12% NuPAGE Bis-Tris gel (Thermo Fisher Scientific). Following electrophoresis, RNA-protein complexes from the gel were transferred onto a nitrocellulose membrane using the Novex wet transfer apparatus. Isotope signals on the membrane were visualized by a Typhoon FLA 9500 (GE Healthcare). Based on the image, the band corresponding to the REFEE-RNA complex (approximately 50 kDa) was excised and transferred to a fresh tube.

Subsequent cDNA library construction steps were conducted according to the protocol of Murano et al.^60^. Libraries were sequenced on an Illumina HiSeq X Ten platform (Macrogen Co., LTD.) with the addition of 30% PhiX control.

### Establishment of Doxycycline-inducible Refee expression in ESCs

The doxycycline-controlled mouse ESC line, TRE::Dux, was established in our previous study^55^. The doxycycline-inducible Refee expression mouse ESC line, TRE::Refee, was generated by co-transfecting EB3 mESCs with pPB-TRE-mVenus-Refee, pPB-CAG-rtTA3G, and pCMV-HyPBase plasmids as described previously^58^. Forty-eight hours after transfection, the cells were subjected to hygromycin (500 μg/ml; FUJIFILM Wako) and G418 (500 μg/ml; FUJIFILM Wako) selection for 7 days. The selected cells were then seeded at 2 × 10^2^ cells/cm^2^ in a culture medium containing hygromycin (250 μg/ml) and G418 (250 μg/ml). Single-cell clones were picked and expanded after 7 days.

### CRISPR/Cas9-mediated gene targeting in ESCs

Plasmids encoding Cas9 and sgRNA against the Refee coding region, pX330-Refee1-Puro and pX330-Refee2-Puro, were constructed in-house. These plasmids were co-transfected into TRE::Dux ESCs using the jet OPTIMUS DNA Transfection Reagent (Polyplus). After 18 hr, cells were selected with 1 μg/mL Puromycin (FUJIFILM Wako) for 54 hr. The selected cells were then seeded at a low density to allow for clonal colony formation. Individual colonies were picked and expanded after 7 days. Expanded colonies were subsequently screened for successful gene targeting by both immunofluorescence analysis of REFEE expression (see “smFISH and Immunofluorescence Analysis” section for details) and genotyping.

For the genotyping, genomic DNA was extracted using the MagExtractor-Genome-kit (Toyobo) according to the manufacturer’s instructions. Approximately 100 ng of genomic DNA was used as template for PCR amplification of regions containing the sgRNA target sites. Primer pairs used were as follows (sequences are listed in Table S8): Cluster 1 *Refee*, primers 361 and Kent2; Cluster 2 *Refee*, primers 052 and 053; *Gm5698*, primers 054 and 055; *Alyref* exon2, primers 162 and 163; *Alyref* exon4, 160 and 161; *Alyref2*, 158 and 159. PCR was performed using KOD Fx DNA polymerase (Toyobo). Thermal cycling conditions were initial denaturation at 94°C for 2 min; 40 cycles of 98°C for 10 s, annealing for 30 s (starting at 60°C and decreasing by 0.2°C per cycle to 52°C), and 68°C for 2 min. Sequencing of the PCR products was performed by the Azenta Sanger sequencing service (Japan).

Cloned Refee KO cells were treated with 2.5 nM Pladienolide B (PlaB, Cayman Chemicals) and/or 1 μg/mL Actinomycin D (FUJIFILM Wako) for 3 days. Following treatment, the cells were harvested for downstream analysis such as quantitative RT-PCR or immunofluorescence.

### Proliferation Assay of ESCs

Proliferation levels of cultured cells were measured using the Cell Counting Kit-8 (Dojindo, Japan) following the manufacturer’s instructions. Briefly, 1.25 ✕ 10^3^ to 5 ✕ 10^3^ cells were seeded into 96-well plates. After overnight culture, PlaB was added to a final concentration of 2.5 nM. Following an additional 2 days of culture, the kit’s solution was added to each well and incubated for 2 hr in a CO_2_ incubator. The colorimetric absorbance pf each well was measured using an iMark microplate reader at 450 nm (Bio-Rad).

### Embryo culture and microinjection

Female mice were superovulated by intraperitoneal administration of 150 μl CARD HyperOva (Kyudo, Japan), followed 48 hr later by 7.5 IU human chorionic gonadotropin (hCG) (Asuka Pharmaceutical, Japan). Subsequently, superovulated female mice were housed with male mice overnight for mating. Successful copulation was confirmed by the presence of a vaginal plug the following morning. Zygotes were collected by dissecting the oviducts at 0.5 days post coitum and were released from cumulus cells using hyaluronidase treatment.

The microinjection was performed as previously described^5^. Briefly, a mixture of anti-Refee ASO #4, #5, and #6 or control ASO (20 uM each, see Table S8 for sequence information) was microinjected into the male pronuclei of the zygotes using a FemtoJet4i microinjector (Eppendorf). Injected embryos were then cultured in KSOM medium (Sigma) at 37 °C with 5% CO2.

### RNA-seq of embryos and ESCs

For embryos, 30 mid-2-cell, late 2-cell, or 4-cell stage embryos were collected per biological replicate. The zona pellucida was removed by treatment with acidic Tyrode’s solution (Merck). Ribosomal RNA-depleted total RNA libraries were prepared using the SMART-seq Stranded Kit (TaKaRa) following the manufacturer’s instructions. Libraries were sequenced on an Illumina HiSeq X Ten platform.

For ESCs, total RNA was extracted using the RNeasy Mini Kit (Qiagen) according to the manufacturer’s protocol. Ribosomal RNA-depleted libraries were prepared and sequenced by Novogene (China).

### Mass spectrometry

TMT-based quantitative mass spectrometry was performed as previously described with several modifications^61^: 100 L2C embryos were collected 24 hr after microinjection and treated with the acidic Tyrode’s solution to remove the zona pellucida. Then the embryos were lysed with 150 μL of guanidine-TCEP buffer (8 M guanidine-HCl (Wako), 100 mM HEPES-NaOH (pH 7.5), 10 mM TCEP (Wako), and 40 mM chloroacetamide (Sigma)). The lysates were dissolved by heating and sonication and cleared by centrifugation at 20,000 *g* for 15 min at 4°C. Proteins were precipitated by methanol/chloroform and solubilized in 50mM triethylammonium bicarbonate buffer containing 0.1% RapiGest SF (Waters). After sonication and vortexing, proteins were digested overnight at 37 °C with 200 ng of trypsin/Lys-C mix (Promega) and were labeled with TMT-10plex reagents (Thermo Fisher Scientific). Labelled peptides were pooled, acidified, and fractionated using offline high-pH reversed-phase chromatography on a Vanquish DUO UHPLC system (Thermo Fisher Scientific). Peptides were separated into 48 fractions, which were consolidated into 16 fractions. Each fraction was evaporated in a SpeedVac concentrator and dissolved in 3% acetonitrile and 0.1% trifluoroacetic acid.

LC-MS/MS analysis was performed on an EASY-nLC 1200 UHPLC coupled to a Q-Exactive Plus mass spectrometer (Thermo Fisher Scientific). Peptides were separated on a C18 reversed-phase column (Nikkyo Technos) using a linear acetonitrile gradient over 180 min. The mass spectrometer operated in data-dependent acquisition mode with top 10 MS/MS. Raw data were searched against the Swiss-Prot mouse database using Proteome Discoverer with Sequest HT, employing standard parameters for trypsin digestion, TMT labelling, and modifications. Quantification was done using reporter ion intensities in Proteome Discoverer software.

The count data from mass spectrometry were analyzed using the limma and sva R packages (v3.64.0 and 3.56.0, respectively). Initial quality control analysis identified four outliers (two from each condition), which were subsequently removed from the set of five samples per condition to reduce inter-sample variability. The specific TMT-labeled samples removed were 126 and 128N (control), and 127N and 131 (*Refee* KD). The remaining sample information, including experimental condition and batch assignment, was used to construct a linear model design matrix. Data were log-transformed and variance-stabilized using the voom method. To address batch effects, we applied ComBat from the sva package, using batch labels and experimental condition as covariates. Negative intensity values resulting from the correction were adjusted by shifting the entire matrix to maintain strictly positive values. Differential protein abundance between *Refee KD* and control samples was assessed using limma’s empirical Bayes framework. Moderated t-statistics were used to rank proteins, and results were exported as a ranked table of differentially abundant proteins. Interactome among differentially expressed proteins was studied and visualized using the STRING web tool (v12.0)^62^.

### smFISH and immunofluorescence analysis

Single-molecule FISH (smFISH) probes against MERVL were designed and synthesized in our previous study^5^ (LGC Biosearch Technologies, Inc., CA). smFISH followed by immunofluorescence staining was performed according to the manufacturer’s protocol. Briefly, Quasar 670-labelled probes were hybridized against MERVL RNA at 37 °C for 16 hr, followed by immunofluorescence staining below.

For immunostaining of embryos, the embryos were fixed in 4% paraformaldehyde for 15 min and permeabilized with 0.5% Triton-X in PBS (Triton-X-PBS), following incubation with the first antibodies at RT. After washing the first antibodies, the secondary antibodies (Thermo Fisher Science, at 1:500 to 1:1000 dilution) and 1 ug/ml DAPI were added to the embryos. The embryos were observed using a Fluoview confocal microscope (FV3000 or FV4000, Evident, Japan) with a UPLSAPO60XS objective. Intensity quantification of the obtained images was analyzed using Fiji (v2.14.0).

For staining culture cells, the cells were fixed in 1% paraformaldehyde for 15 min and permeabilized in Triton-X-PBS. After that, the cells were blocked with 1% skim milk and followed by the protocol of embryo staining. The subsequent samples were observed using a Keyence fluorescent microscope BZ-X1000 (Keyence, Japan). The frequency of marker-positive cells was quantified using the Hybrid Cell Count function in the Keyence BZ-X analyzer software.

### RNA extraction and RT-qPCR

Total RNA was extracted from cultured cells using either ISOGEN (Nippon Gene) or the RNeasy Mini Kit (Qiagen), following the manufacturers’ protocols. Cytoplasmic and nuclear RNAs were extracted from cultured cells using the PARIS Kit (Thermo Fisher Scientific) according to manufacturer’s protocol. Extracted RNA samples were treated with TURBO DNase (Thermo Fisher Scientific) to remove residual genomic DNA.

Reverse transcription was performed using random hexamers and PrimeScript II Reverse Transcriptase (TaKaRa) according to the manufacturer’s instructions. Quantitative PCR was conducted with either TB Green Premix Ex Taq (TaKaRa) or Thunderbird Next SYBR qPCR Mix (Toyobo, Japan) on a Thermal Cycler Dice Real-Time System (TaKaRa). Primers used for this study is described in Table S8.

### iCLIP-seq data processing

Raw single-end iCLIP-seq reads were adapter-trimmed using Cutadapt (v4.3) with the 3′ adapter sequence (TGGAATTCTC) and a minimum read length of 20 bp. Due to abundant adapter concatemers, over 99% of reads had adapter sequences removed. Reads were then collapsed into unique sequences using fastx_collapser (FASTX-Toolkit) to remove PCR duplicates while retaining count information. The first 9 bases, containing barcode and random nucleotides, were subsequently trimmed using fastx_trimmer (-f 10), preserving cDNA-derived sequences for downstream analysis.

Trimmed reads were aligned to the mouse genome (GRCm38/mm10) using STAR (v2.7.4a) in single-pass mode with options --outFilterMultimapNmax 50, -- alignSJoverhangMin 8, and --alignIntronMax 1000000. Sorted BAM files were generated and converted to BED format with bedtools bamtobed. Biological replicates were concatenated and sorted using bedtools sort to prepare input for peak calling.

Peak calling was performed using Piranha (Bioconda), which detects protein-RNA interaction sites by modeling read enrichment. Identified iCLIP peaks were annotated with the ChIPseeker R package using a TxDb object generated from the GENCODE vM25 GTF file via GenomicFeatures. Peaks were annotated relative to nearby genes (±3 kb of transcription start sites) using annotatePeak, with gene symbols assigned via org.Mm.eg.db.

To identify transposable element (TE)-derived peaks, annotated iCLIP peaks were intersected with RepeatMasker-based TE annotations (Dfam mm10_df39) using bedtools intersect with the -wo option to retain detailed overlap information. Only peaks with direct overlap to TE features were retained.

Functional enrichment analysis of identified peaks was conducted using over-representation analysis (ORA) based on hypergeometric tests. Gene ontology (GO) enrichment was performed with the ClusterProfiler R package (v4.16.0).

### RNA-seq data processing

Raw paired-end RNA-seq reads were quality-filtered and adapter-trimmed using Cutadapt with the following parameters: -a AGATCGGAAGAG -A AGATCGGAAGAG -m 20 --pair-filter=any --max-n 0. Trimmed reads were aligned to either the mouse reference genome (GRCm38/mm10) or RepeatMasker-based transposable element (TE) annotations using STAR (v2.7.4a) in two-pass mode with default settings, along with the following additional parameters: --outFilterMultimapNmax 1, --outFilterMismatchNoverReadLmax 0.04, -- alignSJoverhangMin 8, and --alignIntronMax 1000000. Aligned reads were sorted by genomic coordinates.

Gene- and TE-level read counts were computed separately using featureCounts (Subread v2.0.1) with the following options: -T 10 -p -B -C -D 1000 -t exon -g gene_id --fraction -M - O -s 2, and GENCODE mouse annotation (vM25) as reference. Resulting count files were filtered to extract relevant columns (gene ID, read count, and CPM) for downstream analysis. Count data were normalized using edgeR (v4.6.2) in R (v4.5.0).

Gene set enrichment analysis (GSEA, v4.1.0) was performed on ranked gene expression lists. Gene ontology (GO) enrichment analysis was conducted using the ClusterProfiler R package. Splice junction analysis was performed with regtools (v0.5.2) on RNA-seq data from control KD-M2C embryos to identify MERVL-associated chimeric transcripts.

To quantify the genomic proximity of genes to MERVL loci (including MERVL-int and MT2_Mm), gene coordinates (from GENCODE vM25) were first sorted using bedtools sort. The nearest distance to MERVL elements was then computed using bedtools closest with the -d option, which reports the closest feature and the absolute distance between them.

### Data visualization

Three-dimensional structures of proteins obtained from the Uniprot database were visualized and overlaid using Chimera software (v1.13.1). Plots were visualized by ggplot2 (v3.5.2), UpsetR (v1.4.0), ClusterProfiler (v4.16.0), and ggalluvial (v0.12.5) R packages. Read counts, splicing junctions, and Refee iCLIP-seq peaks at the *Aqr* and *Zscan4a* loci were visualized by Integrative Genomics Viewer (IGV, v2.8.0).

### Statistics and reproducibility

All statistical methods, sample sizes, and p-values are described in the corresponding figure legends and/or in the Results section. Briefly, the statistical significance of enrichment analyses (e.g., ORA and GO) was assessed using hypergeometric tests, with Benjamini–Hochberg correction for multiple testing. Differences between two independent groups were evaluated using the Wilcoxon rank-sum test or Student’s t-test, as appropriate. Pearson’s correlation test was used to assess linear relationships between variables. Sequence similarity between GM4340 and ALYREF was evaluated using BLOSUM62-based pairwise alignment. Next-generation sequencing data were generated from two biological replicates, and mass-spectrometry data from three biological replicates.

### Data availability

The RNA-seq and iCLIP-seq data acquired in this study have been deposited in the NCBI Gene Expression Omnibus (GEO) database under accession numbers GSE305444 and GSE305445, respectively. Data are available with an access token, which can be requested from the corresponding author.

## Acknowledgments

We thank Masaru Ariura, Masayuki Sato, Harumi Masuda, Yuka Iwasaki, and Wu Yaning for assisting with the experiments, and Core facility, Collaborative Research Resources, Keio University School of Medicine for the use of their experimental equipment. This work was supported by the MEXT Grant-in-Aid for Scientific Research in Innovative Areas (19H05753 to H.S.), JSPS KAKENHI (25H00008 to H.S., 20K15808, 24K09286, 25H01318, and 2501309 to M.K.I.), Joint Research Programs, Institute of Advanced Medical Sciences, Tokushima University, Medical Research Center Initiative for High-Depth Omics, Takeda Science Foundation (2022049218, M.K.I.), and Keio University Academic Funds (S01JA24199, M.K.I.). This work was also supported by Sumitomo Foundation Research Grant to H.S., and a generous gift from the Medical Corporation Koukoukai to H.S.

## Author contributions

The manuscript was written by M.K.I. and H.S., with critical input from all co-authors. M.K.I., A.S., K.M., and H.S. conceived and designed the study. A.S. performed microinjection to generate *Refee* knockdown embryos and prepared RNA-seq libraries. M.K.I., K.M., Y.G., and T.D.L. generated monoclonal antibodies and *Refee* knockout ESCs and conducted immunostaining experiments. H.I. provided guidance on iCLIP-seq and designed ASOs targeting *Refee*. H.H., T.N., and T.K. maintained the mouse colony and collected embryos. H.K. performed the mass spectrometry analysis. M.K.I. conducted all remaining experiments.

## Supplementary Note

### Supplementary Methods

#### *Refee* cluster alignment

DNA sequences of the *Refee* gene clusters were obtained from the NCBI database and manually trimmed to include only the open reading frame (ORF) regions. Trimmed sequences were aligned by ClustalW (v2.1), and the sequence similarity was assessed with Phylip (v3.697).

#### Phylogenetic tree visualization

Newick-format files were either generated in the above section or downloaded from the Ensembl gene tree. These were visualized using the ggtree R package (v3.16.0).

#### Silver staining

Silver staining of SDS-PAGE gels was performed using the SilverQuest Silver Staining Kit (Thermo Fisher Scientific), following the manufacturer’s protocol.

#### Prediction of RNA-DNA hybrid formation probability

The minimum free energy (ΔG, kcal/mol) for RNA–DNA hybrid formation between guide RNAs and *Refee* genomic sequences, and between antisense oligonucleotides (ASOs) and *Refee* transcripts, was calculated using RNAhybrid (v2.1.2). Sequence alignments were performed using ClustalW, and heatmaps were visualized with the ggplot2 R package.

#### ASO-mediated *Refee* Knockdown in ESCs

*Refee* knockdown ESCs were generated by nucleofection of ASOs targeting *Refee* transcripts (see Table S7 for sequences), using the Lonza Nucleofector IIb system (Lonza). The nucleofection procedure was performed as previously described^5^.

## Supplementary Tables

**Table S1.**

Expression levels and copy numbers of cluster 6 genes at the M2C stage

**Table S2.**

List of *Refee* loci analyzed in this study

**Table S3.**

List of iCLIP peaks identified in this study

**Table S4.**

Differentially expressed genes in *Refee* KO ESC clone #14

**Table S5.**

Differentially expressed genes in *Refee* KD embryos

**Table S6.**

Mass spectrometry data, related to Fig. 5g

**Table S7.**

List of overlapped genes, related to Fig. 6e and Fig. S10a, b.

**Table S8.**

List of oligonucleotides, plasmids, and antibodies used in this study

**Figure S1.**
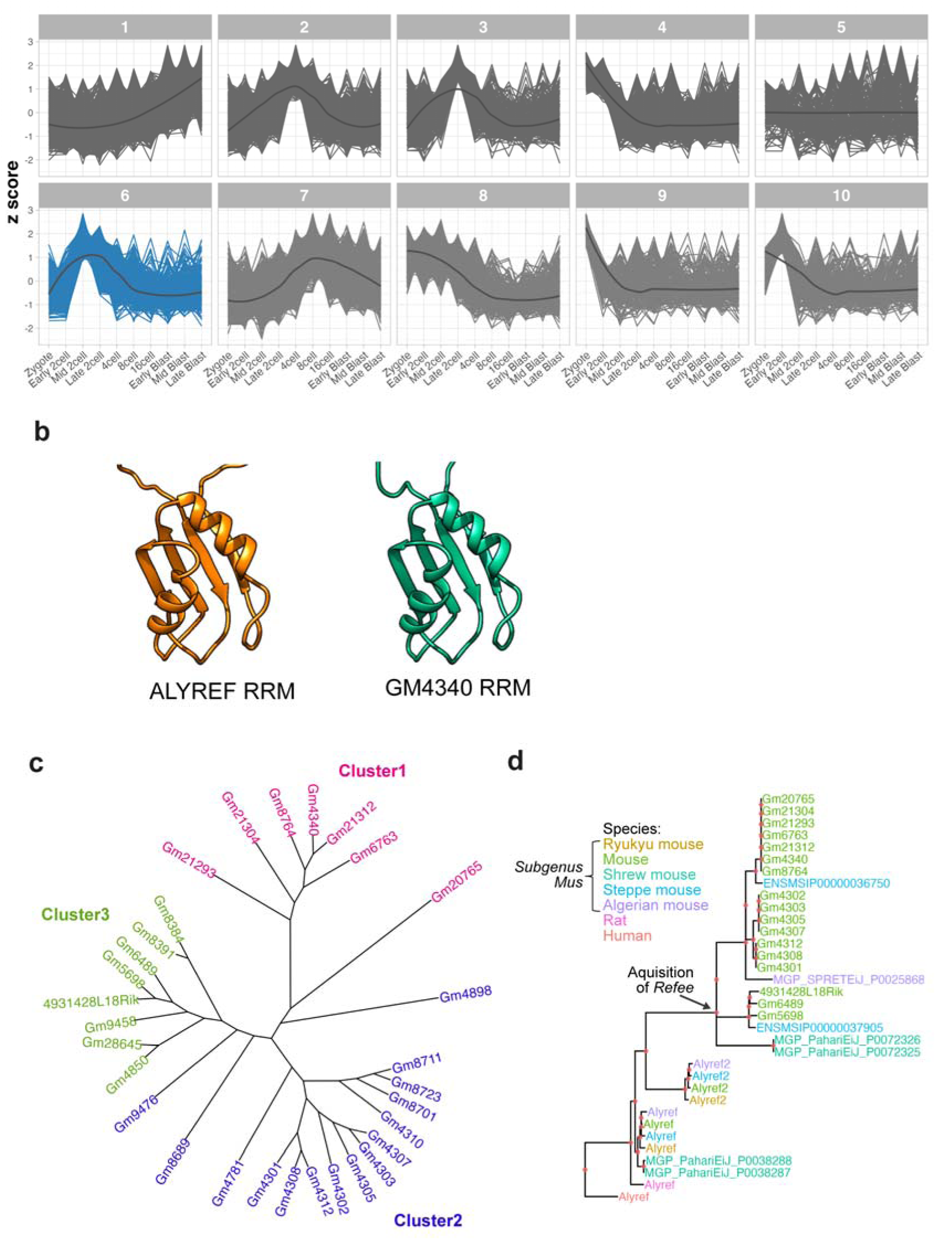
Characterization of *Alyref* family gene expression and evolution. **a.** Expression patterns of 19,760 genes clustered into 10 groups based on z-scores derived from single-cell RNA-seq data by Deng et al.^30^. Blast, Blastocyst **b.** Close-up of predicted RRM conformation of ALYREF (left) and GM4340 (right). **c.** Phylogenetic tree constructed from DNA sequences of the *Refee* gene clusters. **d.** Phylogenetic tree of *Alyref* paralogous proteins among *Mus* subgenus species, rat, and human, showing that *Alyref2* and *Refee* diverged from *Alyref* specifically within the *Mus* subgenus. The data was obtained from the Ensembl gene tree (ENSGT00410000025615).

**Figure S2.**
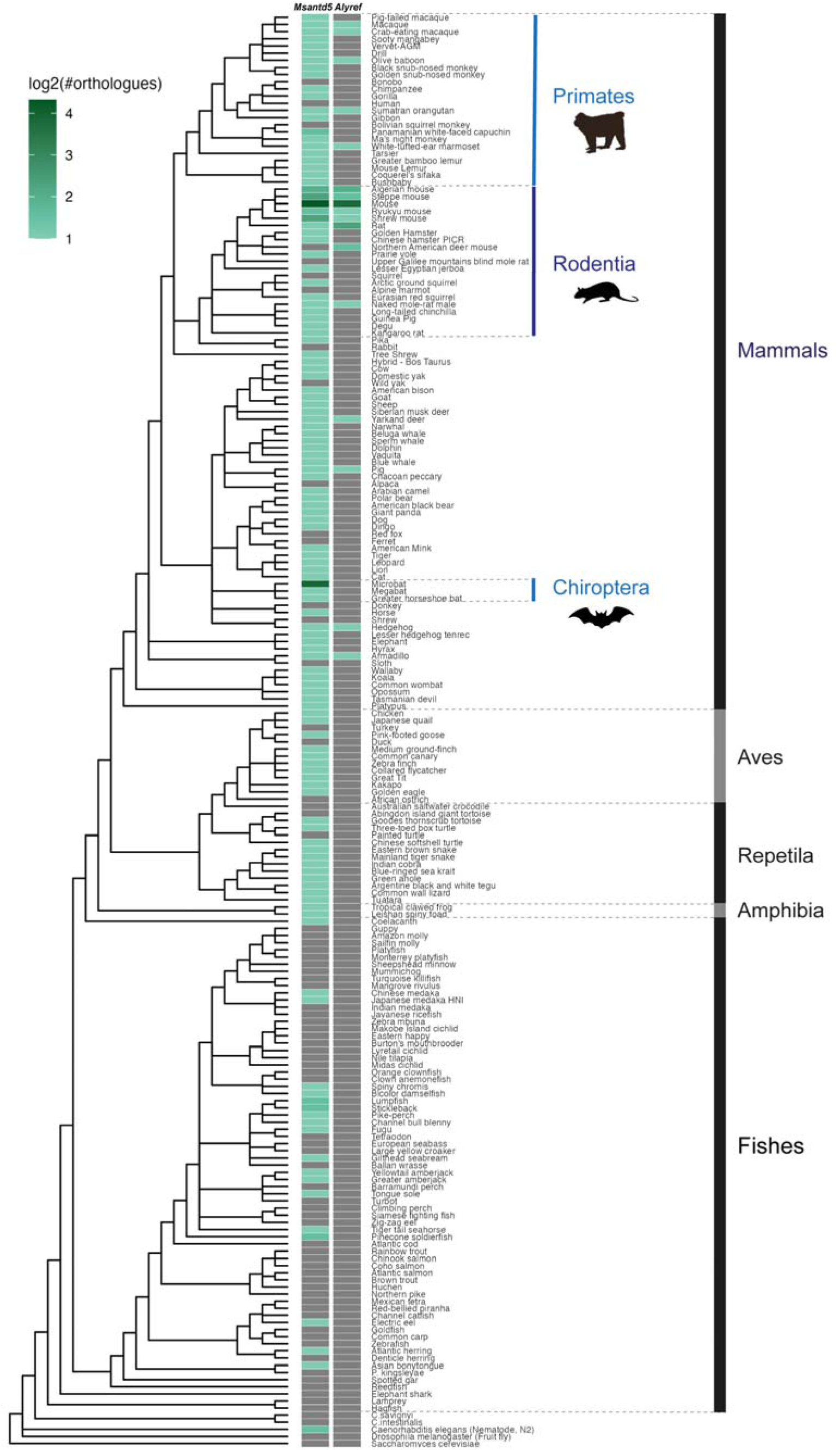
Evolutionary dynamics of *Alyref* and *Msantd5* gene families across species. Heat map combined with a phylogenetic tree, showing the copy numbers of *Msantd5* and *Alyref* genes across representative species. Copy number data were obtained from the Ensembl gene gain/loss tree. Gray tiles indicate the absence of the respective gene in each species.

**Figure S3.**
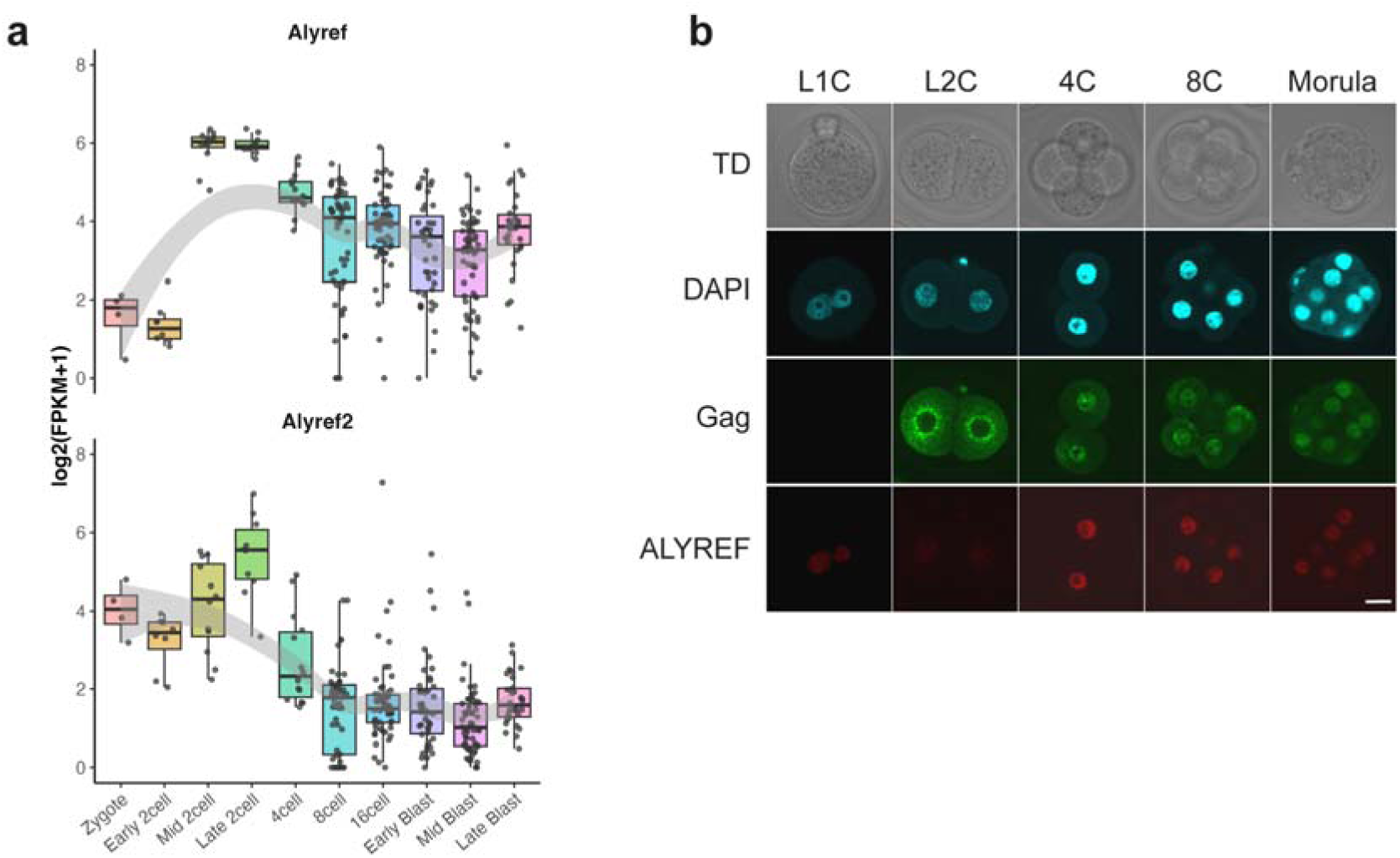
*Alyref* expression during mouse preimplantation embryogenesis. **a.** Expression dynamics of *Alyref* and *Alyref2* during preimplantation development, based on single-cell RNA-seq data from Deng et al.^30^. **b.** Confocal images of zygote to morula stage embryos immunostained for ALYREF. Scale bar: 20µm

**Figure S4.**
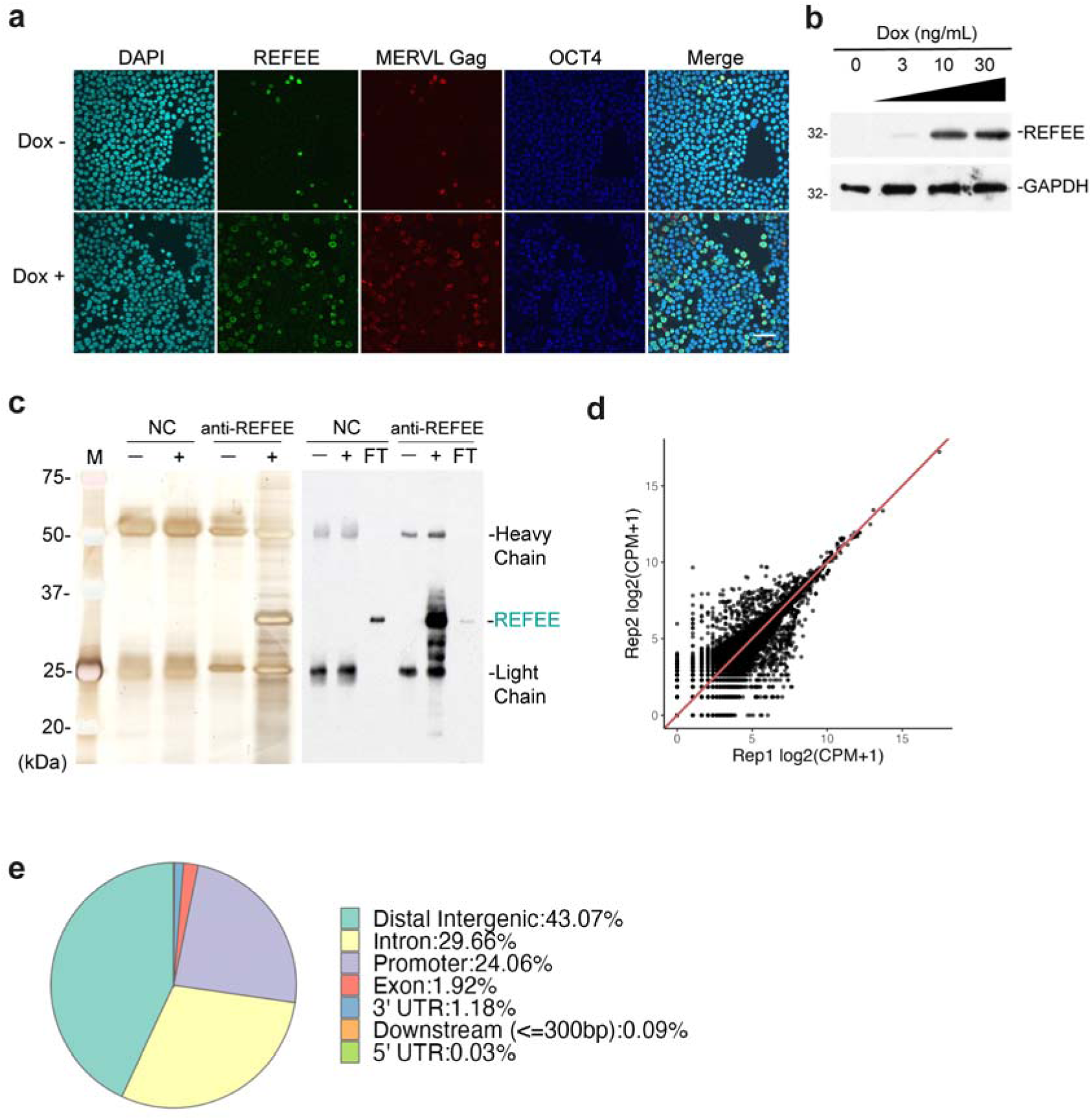
Characterization of REFEE expression and binding in *Dux*-induced 2C- like cells. **a.** Confocal images of doxycycline (Dox)-inducible *Dux*-expressing ESCs (TRE::Dux), with or without Dox treatment. REFEE is expressed in 2CLCs, which are marked by MERVL Gag positivity and loss of OCT3/4. Scale bar: 50 µm. **b.** Western blot illustrating the expression of REFEE protein in TRE::Dux ESCs after Dox-dependent induction of DUX. REFEE expression was analyzed with an anti- REFEE antibody (#8-4). GAPDH level was tested as internal control. **c.** Silver staining (left) and western blot (right) of anti-REFEE immunoprecipitants. + and –; immunoprecipitant with and without cell lysate, respectively, FT; flow- through after immunoprecipitation, NC; Non-Immune IgG, M; Protein molecular weight marker. **d.** Reproducibility of iCLIP-seq across replicates, assessed by Pearson correlation (r = 0.998) **e.** Summary of genomic annotations of REFEE iCLIP-seq peaks.

**Figure S5.**
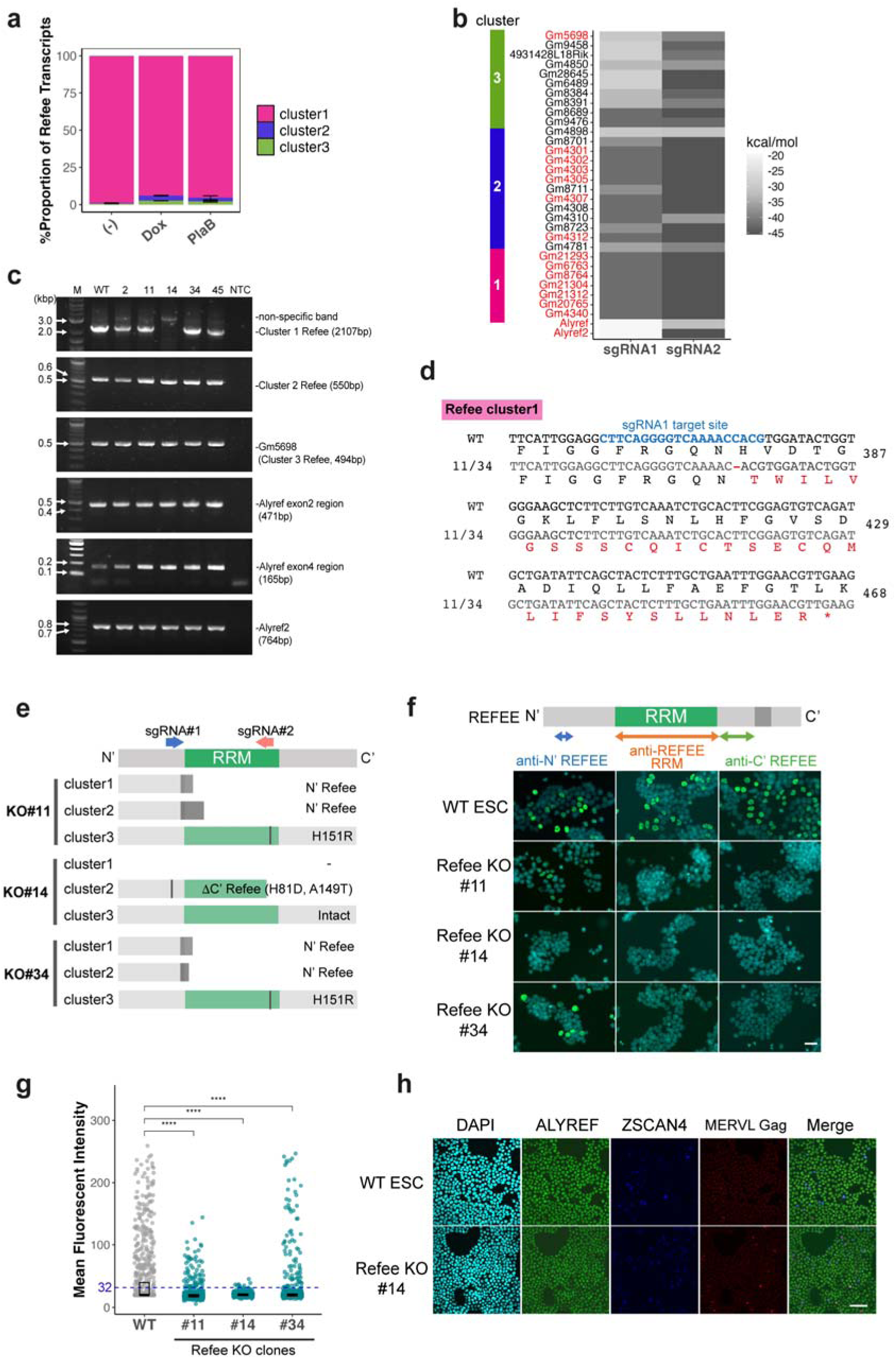
Genetic targeting and expression analysis of REFEE in TRE::Dux ESCs. **a.** Proportion of *Refee* transcripts from each cluster in TRE::Dux ESCs. (-), untreated cells; Dox, cells treated with Dox (10ng/mL) for 24 hours; PlaB, cells treated with Pladienolide B (2.5µM) for 3 days. **b.** Heat map showing the predicted hybridization probability of guide RNAs to *Refee*, *Alyref*, and *Alyref2* genomic regions. The minimum free energy (kcal/mol) was calculated using the RNAhybrid package; lower values indicate higher binding probabilities. Protein-coding genes are highlighted in red. **c.** Genomic PCR analysis using primers specific to each *Refee* cluster, *Alyref*, and *Alyref2*, to evaluate the presence or absence of genomic deletions or mutations. **d.** Genomic sequence of sgRNA1 target site in *Refee* cluster 1. PCR products from panel **c** were sequenced and aligned to the wild-type (WT) sequence (TRE::Dux ESCs). Clones 11 and 34 contain a C-deletion at position 373, resulting in truncated REFEE proteins. **e.** Summary of truncated REFEE variants predicted to be expressed in each knockout ESC clone, based on sequencing data from panel **c**. **f, g**. Immunostaining of REFEE in sgRNA-targeted TRE::Dux ESCs. Panel (**f**) shows representative images stained with antibodies targeting the N-terminus (N′), RNA recognition motif (RRM), and C-terminus (C′) of REFEE. Panel (**g**) provides quantification of REFEE protein expression in individual cells (n = 790) as determined by anti-N′ REFEE antibody staining. The cross bars represent median value ± IQR. Statistical significance is indicated by **** (p < 0.001) using the Wilcoxon rank sum test. Scale bar: 30 µm. **h**. Expression levels of ALYREF in WT and *Refee* KO TRE::Dux ESCs was analyzed by immunofluorescence. Cells were imaged using a confocal microscope. Scale bar: 50 µm.

**Figure S6.**
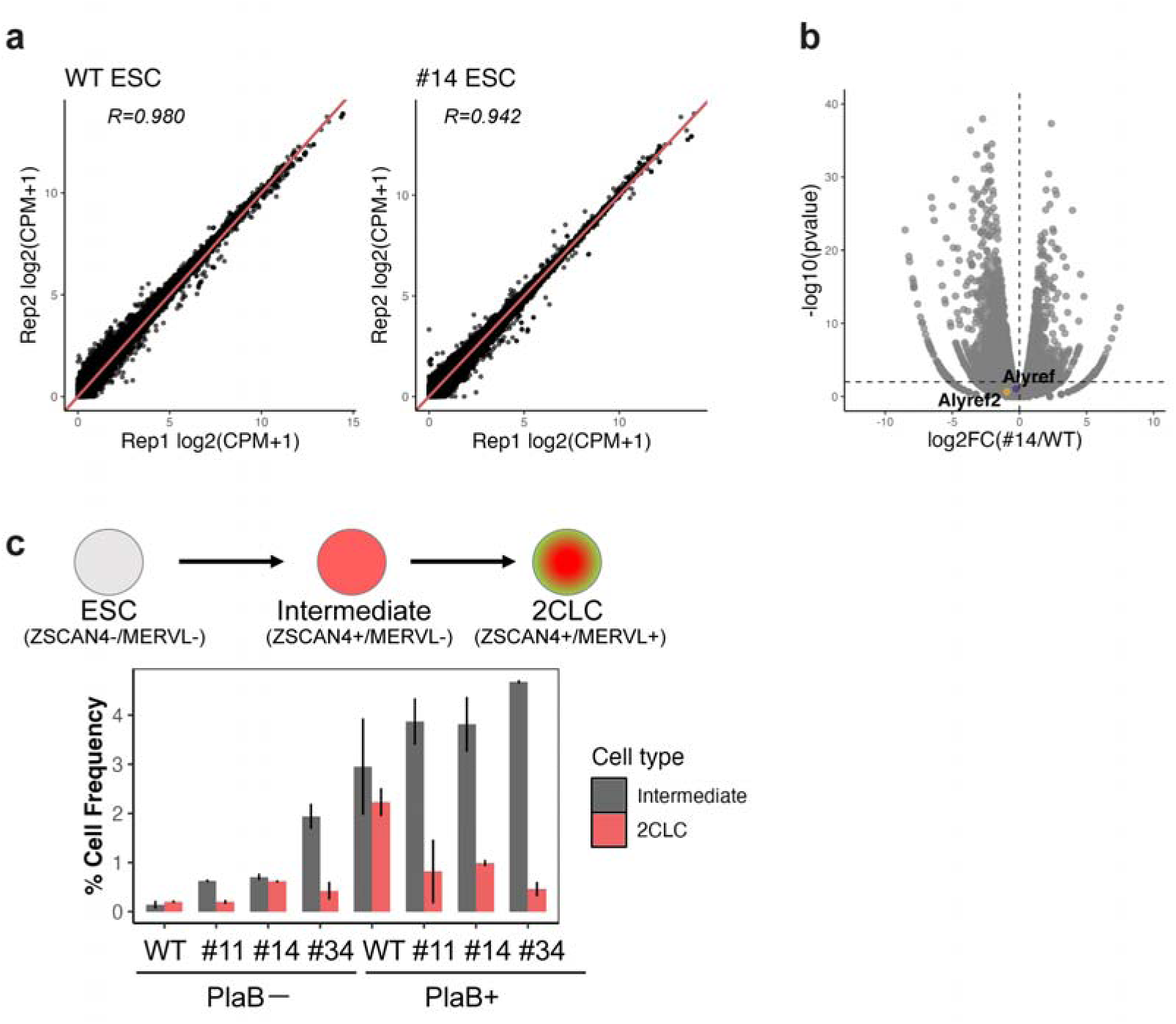
Transcriptome analysis of REFEE KO ESCs and 2CLC conversion efficiency. **a, b** Transcriptome profile of *Refee* KO ESC clone #14 (n=2). (**a**) Correlation of RNA-seq across replicates, showing high reproducibility. R, Pearson’s r. (**b**) Volcano plot demonstrating no significant changes in *Alyref* and *Alyref2* expression in the *Refee* KO clone #14. **c.** Frequency of intermediate (ZSCAN4^+^ single positive cells) and 2CLCs in *Refee* KO ESC clones, with or without PlaB treatment. Quantification was based on two technical replicates (at least 2900 cells per replicate).

**Figure S7.**
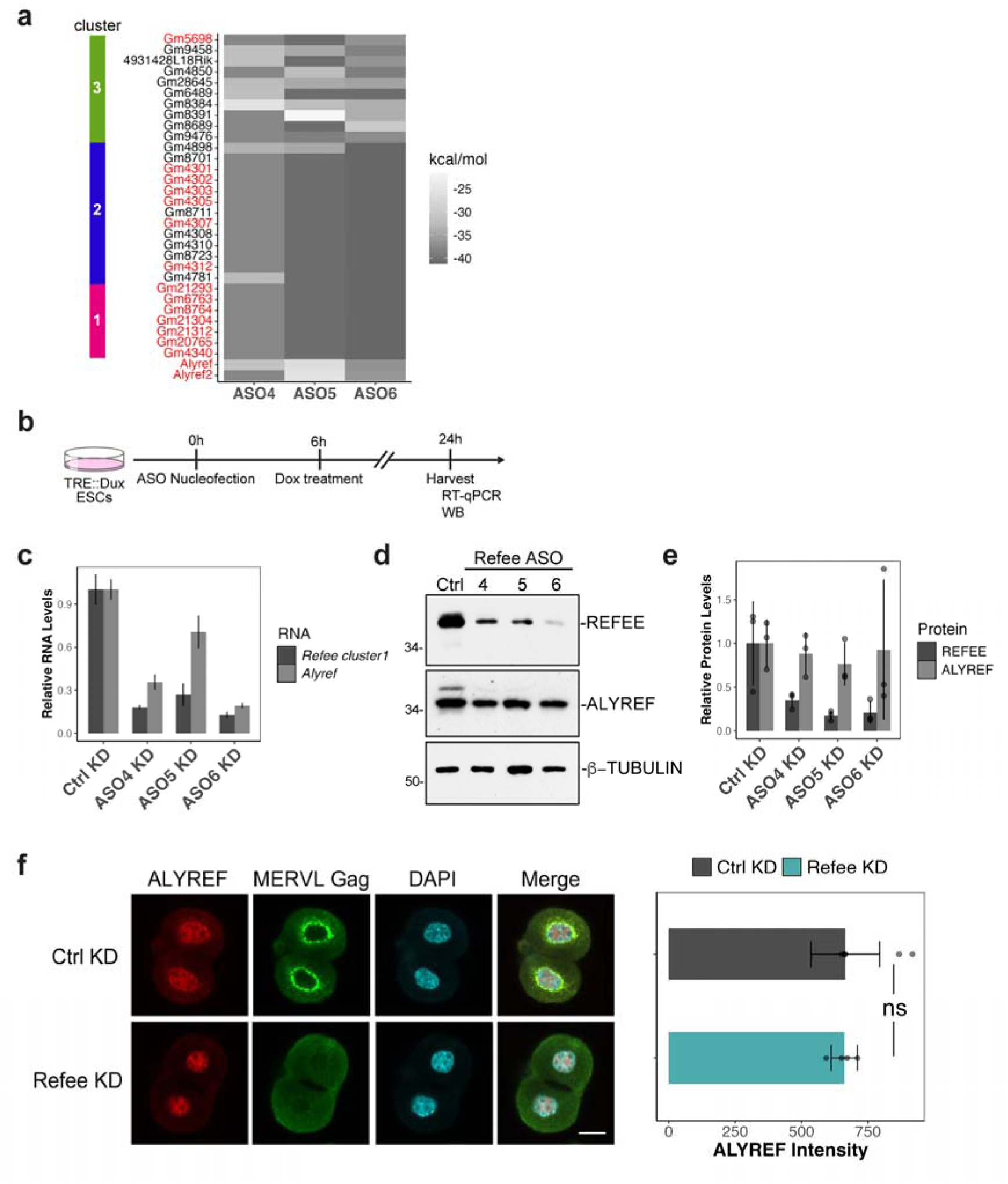
REFEE knockdown efficiency in ESCs and its impact on *Alyref* expression. **a.** Probability of ASO targeting of *Refee*, *Alyref*, and *Alyref2* genomic regions. Heat map of minimum free hybridization energy (kcal/mol), calculated using RNAhybrid. Lower values indicate higher binding probability. Protein-coding genes are shown in red. **b-e.** Knockdown efficiency of *Refee* in ESCs. (**b**) Schematic of the knockdown (KD) experiments. RNA (**c**) and protein (**d and e**) levels of *Refee* and *Alyref* in KD ESCs after 2CLC conversion. Protein levels were quantified based on the obtained images (n = 3) using Fiji (**e**). **f.** Expression of ALYREF in *Refee* KD embryos. Left: Confocal images of L2C embryos immunostained with ALYREF and MERVL Gag. Scale bar: 20µm. Right: Quantification of ALYREF fluorescent intensity per embryo (n = 4).

**Figure S8.**
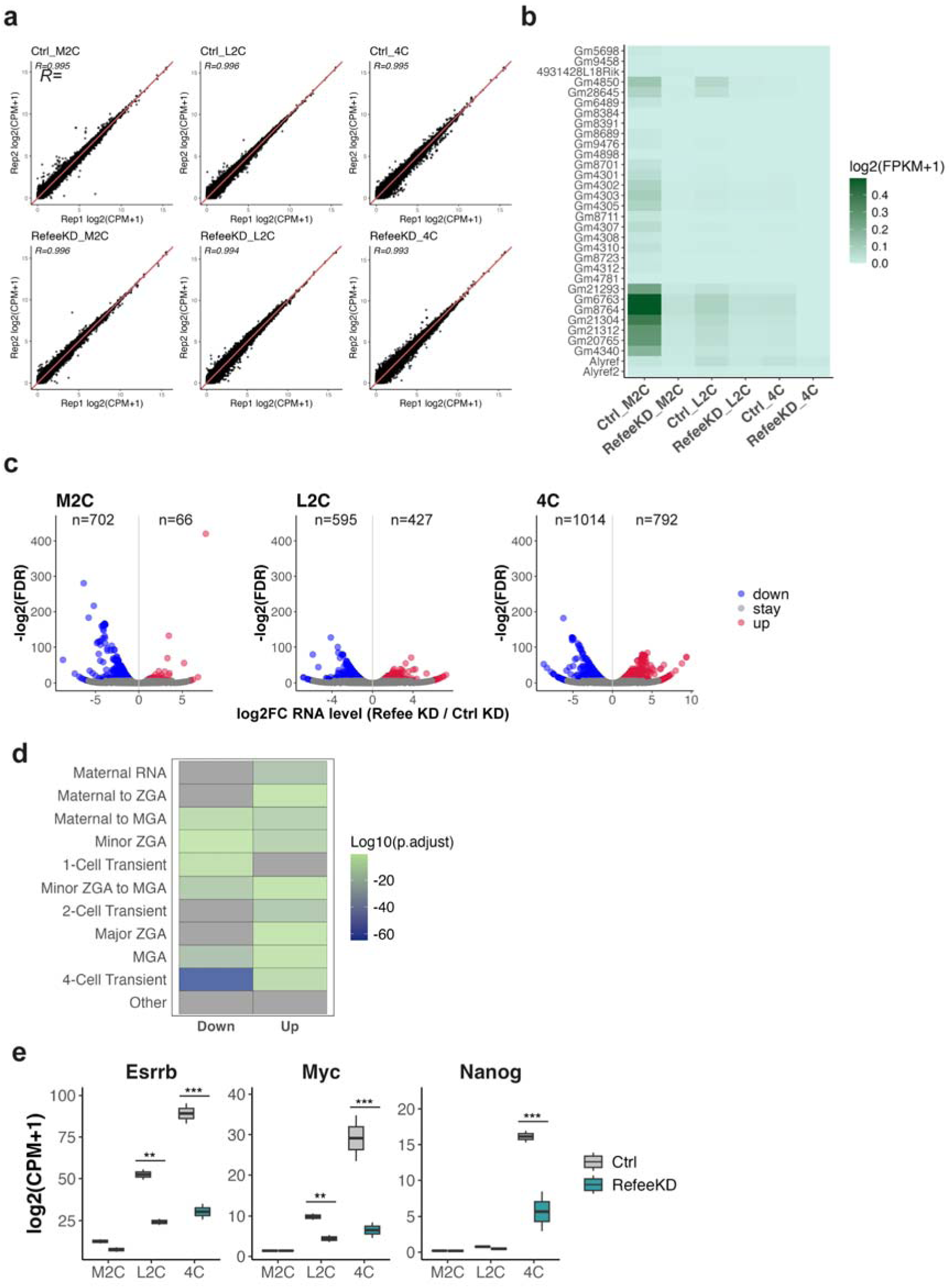
RNA-seq analysis of Refee KD embryos. **a.** Scatter plots showing RNA-seq reproducibility across replicates. Pearson correlation coefficients are indicated. **b.** Heat map showing expression levels of individual *Refee* loci in KD embryos. **c.** Volcano plots displaying differentially expressed genes in *Refee* KD embryos at the indicated developmental stages. **d.** Heat map of gene set enrichment for genes significantly up- or down-regulated in 4C-stage KD embryos. **e.** Expression levels of pluripotency-related genes (*Esrrb*, *Myc*, and *Nanog*) at each stage. Asterisks denote statistical significance (***FDR<0.001; **FDR<0.01).

**Figure S9.**
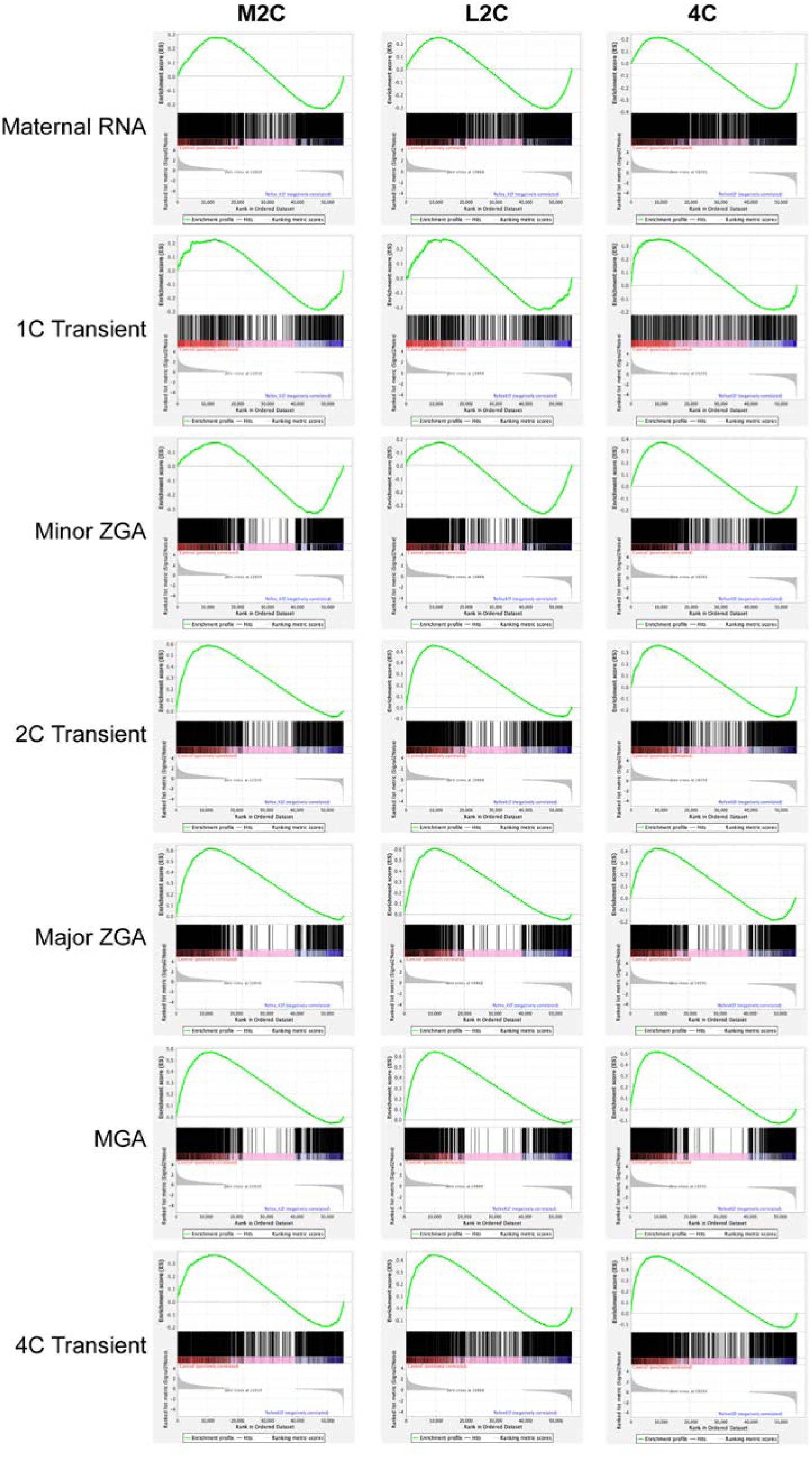
GSEA results related to Fig. 5f. Each panel represents the output of Gene Set Enrichment Analysis (GSEA) for a specific gene set at a given developmental stage. From top to bottom, the plots display: the running enrichment score (ES) curve, the positions of gene set members in the ranked gene list (middle), and the ranking metric scores (bottom), comparing control and *Refee* KD embryos.

**Figure S10.**
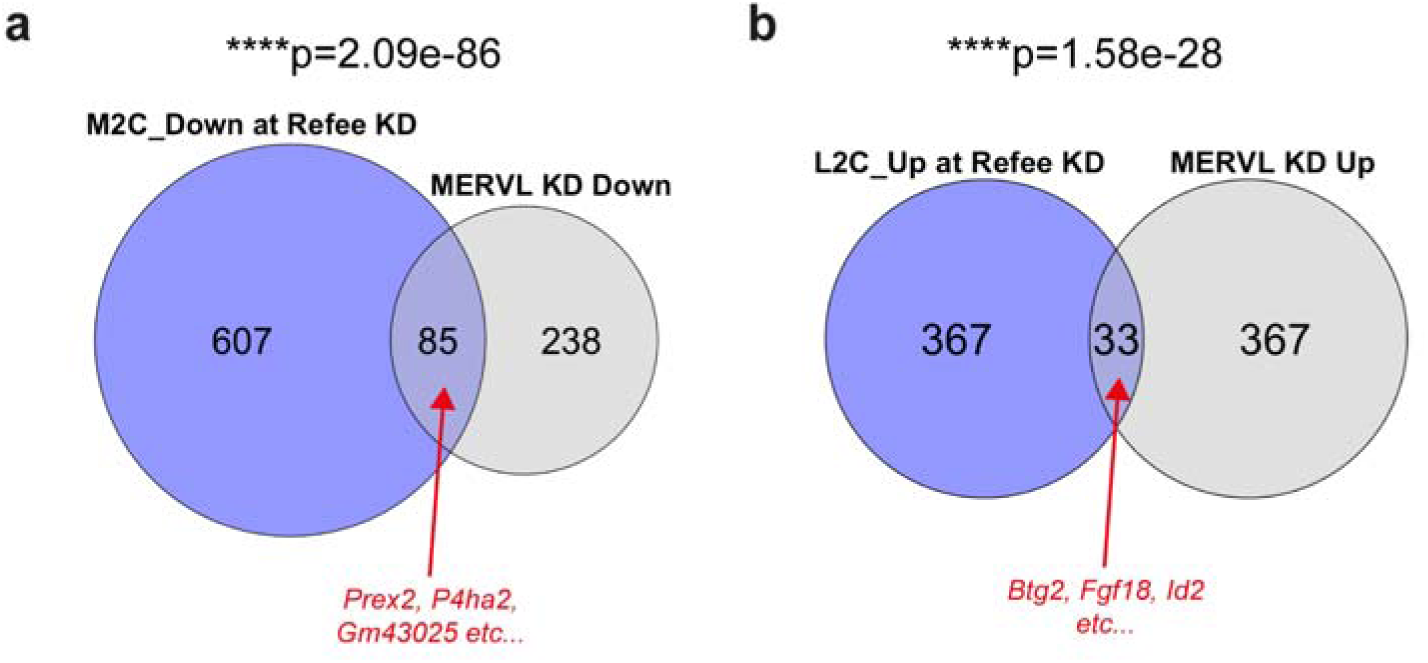
Transcriptomic comparison between *Refee* and MERVL KD embryos. **a, b.** Overlap between M2C_Down genes and downregulated genes in MERVL KD 2C embryos (**a**) and between L2C_Up genes vs upregulated genes in MERVL KD 2C embryos (**b**). Statical significance was calculated by hypergeometric tests.

